# Antibody Correlates of Resilience to *Staphylococcus aureus* Disease and Recurrence in Children

**DOI:** 10.64898/2026.06.26.734609

**Authors:** Mehak Zahoor Khan, Carol M. Kao, Wonyeong Jung, Thendral Selvam, Evgenii Kliuchnikov, Mary G. Boyle, Leslie Fogel, Jeanette Pingel, Jaclyn N. Wright, J. Chase McNeil, Kristina G. Hulten, Sheldon L. Kaplan, Laura Fontana, Doug Lauffenburger, Galit Alter, Juliane Bubeck Wardenburg, Stephanie A. Fritz, Boris Julg

**Author notes:** These authors contributed equally to this work (Mehak Zahoor Khan, Carol M. Kao). Co-corresponding authors, Contact: Boris Julg, MD PhD, Ragon Institute of Mass General, MIT and Harvard, 600 Main Street, Cambridge, MA 02139, USA.

## Abstract

*Staphylococcus aureus* remains a major global pathogen with no licensed vaccine and high recurrent infection burden, yet correlates of protection remain undefined. In a prospective pediatric cohort, we profiled 319 children spanning non-carriers, asymptomatic carriers, those with skin and soft tissue infection (SSTI), or invasive disease. We interrogated 182,149 antibody features, generating the most comprehensive *S. aureus* immune profiling dataset to date. Antibody responses increased with age, marked by expansion of IgG subclasses and Fc-receptor engagement. Asymptomatic carriage was associated with functional antibody profiles targeting conserved surface antigens and select toxins. Multivariate modeling robustly distinguished clinical phenotypes and identified high-value antigens associated with disease resilience. Protection from recurrent disease converged on enhanced FcγR binding and antibody effector function. These findings nominate key antigen targets, and highlight anti-Hla neutralizing antibodies and functional antibodies to additional surface antigens that can be recapitulated through Fc engineering, informing next-generation vaccine and monoclonal antibody strategies.

## Introduction

*Staphylococcus aureus* (*S. aureus)* remains a major contributor to global infectious disease morbidity and mortality^1^. As an opportunistic pathogen, it causes a broad spectrum of clinical manifestations ranging from superficial skin and soft tissue infections (SSTIs) to life-threatening invasive infection and bacteremia^2^. In the United States, invasive *S. aureus* infections result in more deaths annually than any other single infectious agent, surpassing the number of deaths associated with HIV/AIDS and viral hepatitis combined.^.^ More than one in four Americans seek treatment for SSTI each year and up to 70% of individuals experience recurrent infections^3^. The emergence of community-acquired methicillin-resistant *S. aureus* (CA-MRSA) strains has transformed *S. aureus* from a nosocomial concern into a widespread community threat^4^, further amplifying the global burden of antimicrobial resistance and underscoring the urgent need for novel preventive strategies.

Despite decades of research and over 35 clinical trials, no vaccine or antibody-based therapy against *S. aureus* has been licensed. Previous vaccine efforts have focused heavily on a limited set of canonical antigens, including clumping factors (ClfA/ClfB), iron-regulated surface determinants (IsdA/IsdB), α-toxin (Hla), and capsular polysaccharides^5, 6, 7^. Although these antigens conferred robust protection in animal models^8, 9, 10, 11, 12, 13, 14, 15, 16, 17, 18^, they failed to elicit protective immunity in humans. A growing body of literature suggests that these failures are not due to the inability of the host to generate an anti-staphylococcal immune response. Indeed, humans are exposed to *S. aureus* in the first months of life, exhibiting diverse antibody responses against staphylococcal antigens as a function of carrier state^19, 20, 21^. Vaccine failures are thus proposed to stem from a combination of pre-exisiting imprinted non-protective responses, *S. aureus* immunoevasion, virulence factor diversity and redundancy, and the lack of identification of antibody specificities that are functionally protective in individual hosts^22^. These setbacks underscore the urgent need to deepen our mechanistic understanding of protective immunity and utilize this knowledge to revisit vaccine design.

While higher infection rates and poorer outcomes with *S. aureus* are observed in immunocompromised populations, including individuals with primary immunodeficiency such as chronic granulomatous disease, or aquired immunodeficiency as in HIV, cancer, and transplant recipients^23, 24, 25, 26, 27, 28, 29^, severe and recurrent *S. aureus* disease is disproportionately higher very early in life and in older adults^30^. These data suggest that investigation of the host immune response to *S. aureus* as it develops in childhood may provide insight on the biological signatures of relative resilience against *S. aureus* disease. Linked to emerging data that poor control of *S.aureus* may be related to the evolution of functionally impaired *S.aureus* humoral immune responses either due to hyper Fc-sialylation that dampens functional humoral activity as has been observed for IsdB^31^, or through the induction of suboptimal activity as observed to Hla neutralization^32^, here we aimed to examine the humoral immune profiles across young children with differential control of *S.aureus* to capture natural resilience to the bacteria.

We hypothesized that resilience will reflect a more functional antibody profile to particular *S.aureus* antigens, consistent with the observation that antibodies from children recovering from invasive *S. aureus* infection protect mice against lethal challenge^33^, and monoclonal antibodies (mAbs) targeting ClfB, Hla, IsdB, LukAB, and SpA^17, 34, 35, 36, 37, 38^ confer protection in animal models. To test this, we profiled antibody responses in a unique prospective observational pediatric cohort, the PISA Study (Pediatric Immunity to *Staphylococcus aureus*). Using a comprehensive, high-dimensional antibody functional profiling platform (Systems Serology^39^), we characterized humoral responses across children with distinct *S. aureus* disease phenotypes and levels of protection from reinfection. Machine learning approaches were then applied to define the antibody features that most strongly correlate with protection from *S. aureus* disease.

## Results

### Antibody profiling distinguishes colonization from disease in pediatric *S. aureus* infection

We systematically charted the functional humoral immune response to *S. aureus* in a pristinely characterized pediatric cohort called the PISA Study (*Pediatric Immunity to Staphylococcus aureus*) spanning early infancy though young adulthood (age range 0.04-20.35 years) and balanced by sex (160 male and 159 female). The study comprises four distinct clinical groups: (i) healthy controls without *S. aureus* carriage at the time of enrollment, (ii) healthy controls with *S. aureus* carriage at enrollment, (iii) subjects with culture-confirmed *S. aureus* SSTIs (e.g. abscesses, cellulitis), and (iv) subjects with invasive *S. aureus* infection (such as bacteremia, pneumonia, or osteoarticular infection) **(Table S1 and S2)**. Across a subset of these subjects **(Figure 1a)**, we utilized a custom Luminex systems serology platform^39^ to profile humoral immune responses in a blinded manner against 27 *S. aureus* antigens encompassing toxins, immune evasion, surface proteins, and 5 control antigens **(Table S3)**, measuring IgG, IgG1-4, IgA1-2 and IgM responses and FcγR1-4/FcαR binding at acute (V1) and convalescent (V2) time points (6-8 weeks after acute infection) **(Table S4)**.

**Figure 1.**
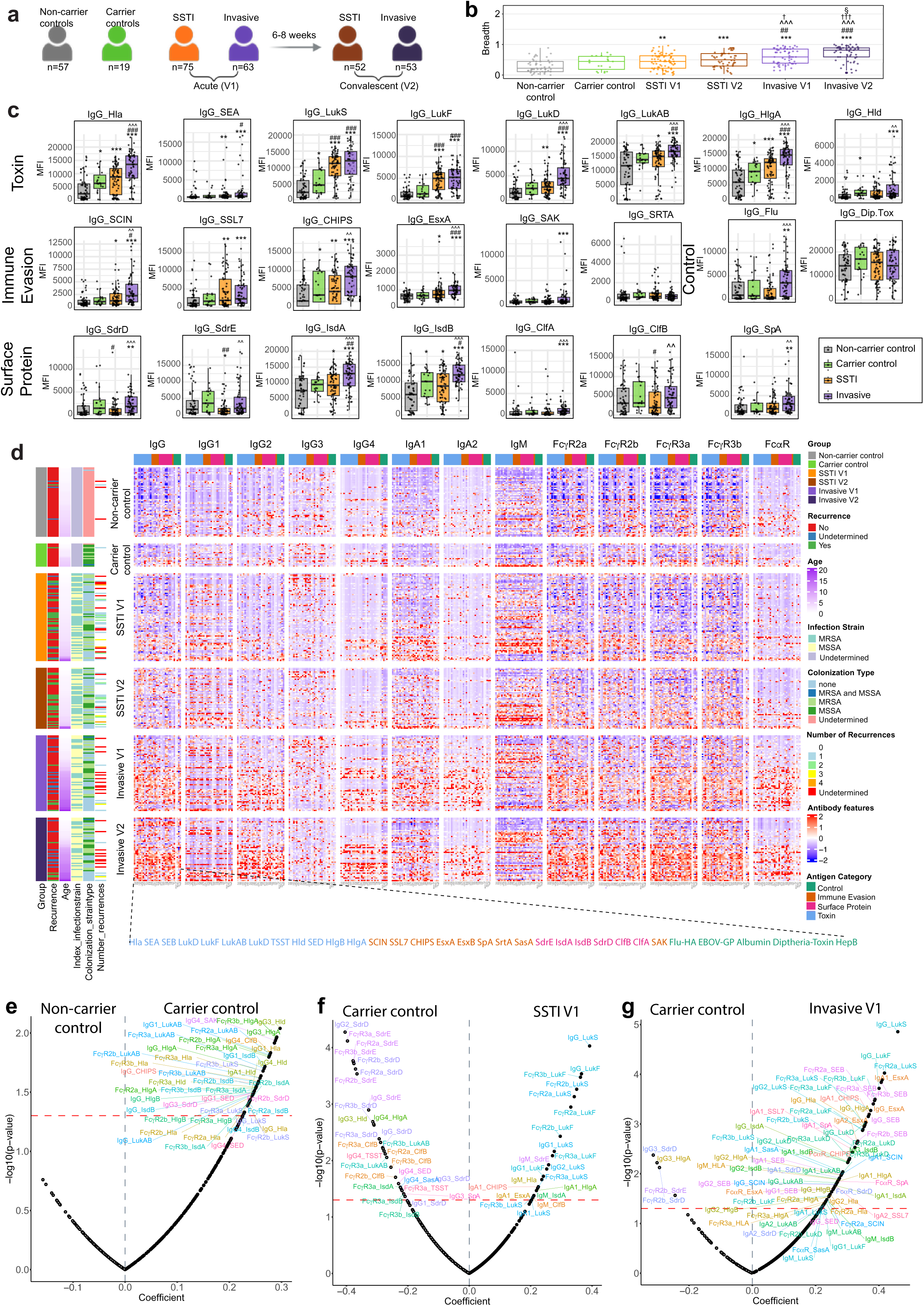
Colonization drives focused, opsonic antibody responses while SSTI/invasive disease triggers broad inflammatory responses. (a) Schematic overview of the Pediatric Immunity to *Staphylococcus aureus* (PISA) cohort design. Children from birth to 21 years of age were stratified into healthy non-carrier control or carrier (asymptomatically colonized) control, SSTI (skin and soft tissue infection), and invasive disease groups based on clinical presentation and culture-confirmed *S. aureus* infection. Serum samples were collected at acute (V1) and convalescent (V2) timepoints and profiled using a customized Luminex-based systems serology platform. (b) Boxplots highlighting the interquartile range (IQR) and median show the antigen breadth (fraction of staphylococcal antigens recognized) in serum IgG across six groups: non-carrier control, carrier control, SSTI patients at Visit 1 (V1) and Visit 2 (V2), and Invasive infection patients at V1 and V2. Each dot represents an individual. Statistical comparisons between groups were performed using the Mann–Whitney U test, with horizontal bars indicating pairwise significance: **p < 0.01, ***p < 0.001 comparisons to healthy non-carrier controls; ##p < 0.01, ###p < 0.001 indicate comparisons to carrier individuals, ^^^p < 0.001 indicate comparisons to SSTI V1, † p < 0.05 indicate comparisons to SSTI V2 and §p < 0.05 indicate comparisons to Invasive V1 (c) Antigen-specific IgG responses to *S. aureus* antigens across four groups: non-carrier control (gray), carrier control (green), SSTI V1 (orange), and invasive V1 (purple). Antigens are grouped by functional category: toxins (top row), immune evasion factors and control (middle row), and surface proteins (bottom row). Each point represents an individual; box plots show median and IQR. Statistical comparisons between groups were performed using the Mann–Whitney U test, with stars indicating pairwise significance (*p < 0.05, **p < 0.01, ***p < 0.001) comparisons to healthy non-carrier controls; hashes indicate comparisons to carrier individuals (#p < 0.05, ##p < 0.01, ###p < 0.001), and carets indicate comparisons to SSTI (^p < 0.05, ^^p < 0.01, ^^^p < 0.001). (d) Heatmap of z-scored antibody responses (rows = samples, columns = antibody features) across all subjects. Features include total IgG, IgG1–4, IgA1/2, IgM, and Fc receptor binding (FcγR2a, FcγR2b, FcγR3a, FcγR3b, FcαR) against all antigens. Samples are grouped by clinical category, sorted by age within each clinical category and annotated for recurrence, infection strain, colonization strain type, and recurrence history. Antigen classes are color-coded (green = control, orange = immune evasion, pink = surface protein, blue = toxin). (e-g) Volcano plots showing univariate comparisons between groups. Each point represents an antibody feature plotted by effect size (x-axis: regression coefficient) and significance (y-axis: –log10 p-value). Features significantly enriched in non-carrier vs. carrier control (e), SSTI V1 (f), or invasive V1 disease (g) are labeled. The horizontal red dashed line marks significance (p = 0.05). Features are colored by antigen specificity.

To quantify the overall humoral response to *S.aureus*, we calculated the fraction of staphylococcal antigens for which each child exhibited an above-median IgG response across the six study groups **(Figure 1a)**. The breadth of antigen recognition increased progressively from healthy non-carriers to carriers, rising further in SSTI patients, with the highest observed in individuals with invasive infections, suggesting a strong association between disease severity and the breadth of response. Moreover, breadth also increased from V1 to V2 within populations both with SSTI and invasive infections, reflecting a broadening of the IgG response over time. Importantly, all healthy controls displayed detectable IgG responses, consistent with exposure to *S. aureus*^19, 20, 21^. IgG responses to most toxins scaled with disease severity, peaking in children with invasive infections. In contrast, differences in IgG levels to immune evasion antigens were more moderate, though elevated responses were still observed in both SSTI and invasive cases compared to healthy non-carrier and carrier controls (**Figure 1b**). Strikingly, several surface protein antigens, such as ClfB, IsdB, SdrD, and SdrE, elicited significantly higher IgG responses in healthy *S. aureus* carriers compared to SSTI cases, potentially indicating protective immune signatures **(Figure 1c).**

Systems-level heatmap visualization highlighted that beyond IgG, other subclasses, isotypes and FcR binding levels were markedly reduced in the non-carrier controls **(Figure 1d).** Invasive cases, particularly at V2, exhibited the strongest and broadest antibody responses across nearly all antibody features. SSTI cases exhibited a more moderate profile dominated by IgM, while healthy children with *S. aureus* carriage showed selective enrichment of antibodies to specific surface proteins. Importantly, IgG (except IgG3) and IgA subclasses and FcR binding increased with age within each disease group **(Figure 1d, Figure S2).**

Next, to identify markers associated with distinct clinical outcomes, we compared well-defined clinical states. Healthy carriers displayed a unique antibody profile, characterized by elevated IgG, IgG1, IgG3 and FcγR binding responses targeting toxins (Hla, Hld, HlgA/B, LukAB, LukS, SED), surface proteins (ClfB, IsdA/B, SdrD), and immune evasion factors (CHIPS, SAK). These features were absent in non-carriers, suggesting that colonization—though clinically silent—is immunologically active, driving subclass-switched responses, likely through mucosal priming **(Figure 1e)**. Compared to SSTI cases, all healthy children demonstrated selective enrichment of opsonic IgG and FcγR binding to ClfB, IsdA/B, and SdrD/E proteins and other select surface and toxin proteins. In contrast, SSTI cases showed broad IgG, IgM and FcγR binding responses directed to cytolytic toxins indicative of an acute-phase, inflammatory humoral response **(Figure 1f, S1a, S1c-e)**. This contrast became even more striking when comparing healthy carriers with children who had invasive disease. The latter exhibited high-magnitude, multi-isotype responses spanning a broad antigenic spectrum, whereas the former showed selective enhancement of IgG3 and FcγR binding to SdrD/SdrE, along with IgG3-HlgA **(Figure 1g, S1b, S1h)**. These data suggest that invasive disease is characterized by broad, inflammatory, and toxin-focused humoral profiles, whereas healthy carriers possess distinct functionally mature antibody signatures that are more focused and potentially protective **(Figure S1i-j)**.

Age distribution analysis revealed that children with invasive disease were older **(Figure S2a)**, consistent with the known epidemiology of *S. aureus* disease in the pediatric population. Importantly, most antibody features exhibited strong positive correlations with age, consistent with cumulative immune exposure and maturation of humoral responses over time **(Figure S2b, S3)**. Paired analysis of IgG responses from SSTI patients and children with invasive disease from the acute and convalescent stages revealed significant heterogenity in antibody titers **(Figure S4)**. After adjusting for age, healthy carriers retained higher IgG and FcγR binding to surface adhesins (ClfB, IsdA/B, SdrD/E) and select toxins (Hld, LukAB, TSST), while SSTI and invasive infections remained dominated by LukS/LukF responses **(Figure S5a-i)**. A four-way comparison identified ClfB, HlgA, SdrD, and SdrE, as antigens consistently enriched in healthy colonized individuals **(Figure S5j)**. Collectively, these findings indicate that *S. aureus* carriage during childhood elicits a durable, surface-focused opsonic antibody profile, distinct from the toxin-biased responses observed during clinical disease.

### Functional antibody profiling reveals distinct qualitative signatures across clinical states

While elevated antibody titers are a hallmark of infection, emerging evidence suggests that antibody quality plays a more decisive role in protection than the total magnitude of the response alone^31, 40, 41,22,23^. Hence, we next examined a suite of functional metrics: (i) IgG1 avidity index, (ii) lectin-binding glycosylation profiles, quantified by Ricinus communis agglutinin I (RCA) and Sambucus nigra agglutinin (SNA) binding, which serve as surrogates for galactosylation of the IgG Fc domain and terminal sialylation^42^, respectively, and (iii) Fc effector functions -antibody-dependent-complement deposition (ADCD), neutrophil phagocytosis (ADNP), and cellular (monocytic) phagocytosis (ADCP). While lectin-binding antibody features were found to be age-independent, all other metrics were age-corrected to isolate disease-specific features **(Figure S6)**.

IgG1 avidity, a surrogate for affinity maturation, was lowest in healthy children across antigens (except SrtA), irrespective of carriage state **(Figure 2a)**. In contrast, individuals with SSTI and invasive infection —particularly at convalescent timepoints—showed significantly elevated avidity to multiple antigens, including IsdA, IsdB, and several leukotoxins. Notably, IgG1 responses to EsxB, SAK, SED, SasA, and SpA possessed low avidity across all groups, pointing to either limited immunogenicity or effective immune evasion by these antigens. SSL7-specific antibodies were markedly less avid in healthy carriers. EsxA-specific antibodies reached peak avidity in children with SSTI, while Hla-specific antibodies were most avid in healthy carriers and invasive cases. Interestingly, ClfB and SEA antibodies achieved their highest avidity in convalescent samples, indicating that prolonged antigen exposure drives maturation against select surface proteins and toxins, indicative of ongoing immune maturation post-infection.

**Figure 2.**
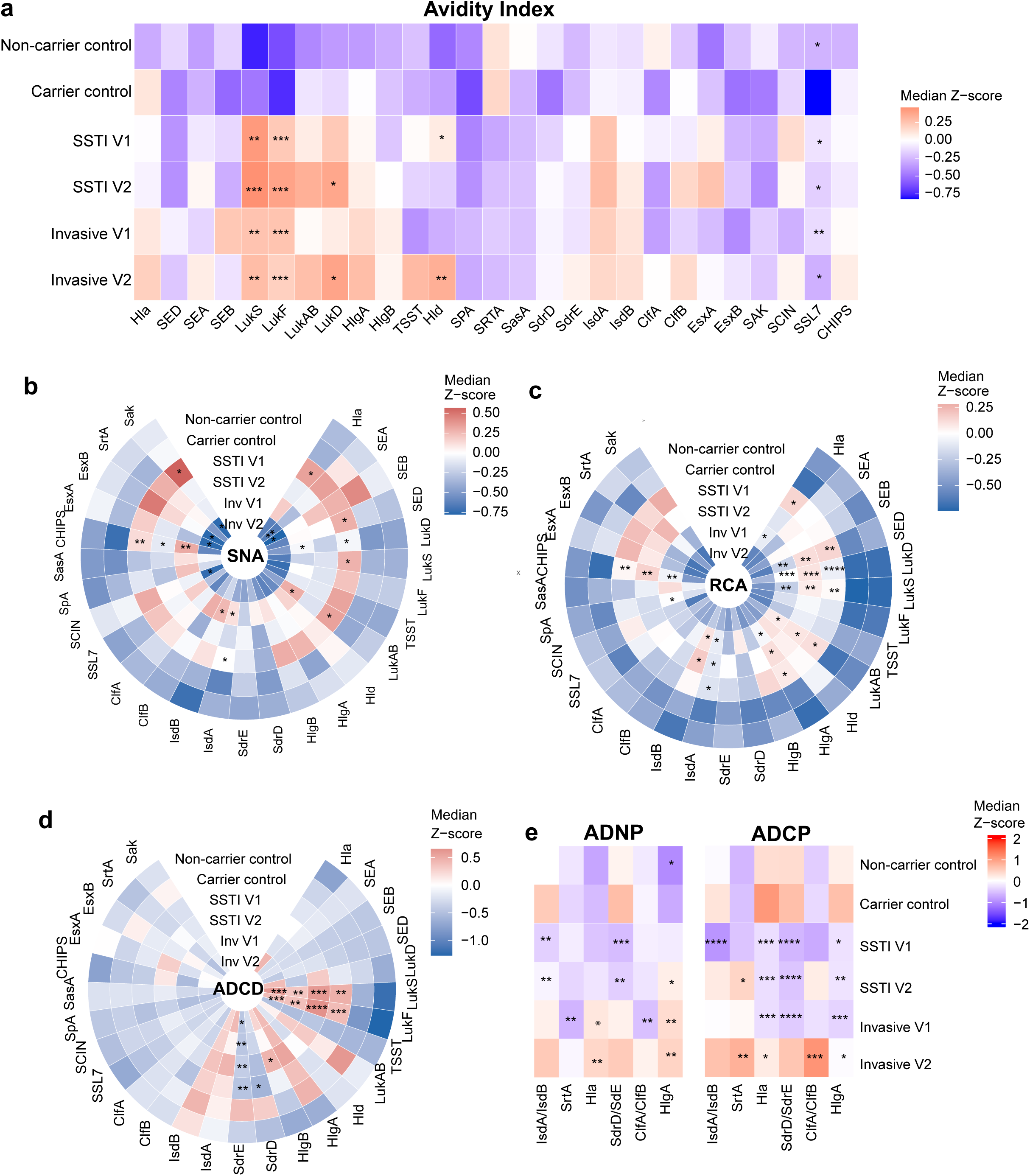
Functional and Glycosylation Profiling Reveals Qualitative Differences in Antibody Responses Across Clinical Groups. (a) IgG1 avidity was quantified using a urea-elution Luminex assay, wherein antigen-specific IgG1 levels were measured in the presence and absence of 7 M urea. Avidity index was calculated as the ratio of MFI with urea to MFI without urea, reflecting the strength of antigen–antibody interaction. Individual avidity scores were age-corrected by residuals and Z-scored for each antigen-specific response. Median Z-scores of each clinical group is shown, with higher Z-scores (red) indicating increased relative avidity and lower Z-scores (blue) indicating reduced avidity. Statistical comparisons were performed between each group and the carrier control using Mann–Whitney U tests (*p < 0.05, **p < 0.01, ***p < 0.001). (b-c) Radial heatmap depict lectin profiling of antigen-specific antibodies. (b) SNA binding serves as a surrogate for Fc sialylation, and (c) RCA binding serves as a proxy for Fc galactosylation. Heatmaps display median Z-scores per antigen-specific feature, with red and blue representing higher and lower lectin binding, respectively. Statistical comparisons were performed between each clinical group and the carrier control using the Mann–Whitney U test with Benjamini–Hochberg (BH) correction. Asterisks denote adjusted p-values (*p ≤ 0.05, **p ≤ 0.01, ***p ≤ 0.001, ****p ≤ 0.0001). (d) Radial heatmap showing median Z-scored antibody-dependent complement deposition (ADCD) against each *S. aureus* antigen across clinical groups. MFI values were age-corrected by residuals and median Z-scores were calculated for each antigen per clinical group. Red and blue indicate higher and lower ADCD activity, respectively, relative to the healthy carrier control. Statistical comparisons were performed using the Mann–Whitney U test with BH correction (*p ≤ 0.05, **p ≤ 0.01, ***p ≤ 0.001). (e) Heatmap showing median Z-scored antibody-dependent neutrophil phagocytosis (ADNP) and antibody-dependent cellular phagocytosis (ADCP) against each *S. aureus* antigen across clinical groups. Phagocytic scores were age-corrected by residuals and Z-scored across each antigen. Heatmap demonstrates median Z-scores for each clinical group, with red and blue indicating increased and decreased ADNP and ADCP activity, respectively, relative to the healthy carrier control. Statistical comparisons were performed using the Mann–Whitney U test with BH correction (*p ≤ 0.05, **p ≤ 0.01, ***p ≤ 0.001).

We next assessed Fc glycosylation profiles using SNA and RCA binding analysis **(Figure 2b-c)**. Healthy children exhibited the lowest lectin binding irrespective of carrier status. By contrast, acute SSTI and invasive infections were associated with sharp increases in both SNA and RCA binding across multiple antigens, including CHIPS, Hla, HlgA, IsdB, LukAB, LukD, LukF, LukS, and SED, indicating a shift toward Fc glycoforms with reduced functional capacity during active disease. Notably, SNA binding normalized at convalescent timepoints, whereas RCA binding remained elevated during convalescence in the SSTI group, suggesting differential glycosylation remodeling over time. Together, these data imply that children who resist infection maintain a more functionally protective glycan profile.

To determine whether enriched responses to specific antigens in healthy carrier individuals represent biomarkers or mechanistically contribute to disease resistance, we prioritized six antigens consistently elevated in healthy children versus disease groups for targeted functional analysis: ClfA/ClfB, Hla, HlgA, IsdA/IsdB, SdrD/SdrE, and SrtA **(Figure 2d-e)**. Individuals with SSTI and invasive infections exhibited robust antibody-dependent complement deposition (ADCD) responses against leukotoxins LukF and LukS at both timepoints. Notably, SdrD-and SdrE-specific ADCD activity was highest in healthy carriers. In contrast, IsdA-, LukAB-, and HlgA-specific ADCD activity was absent only in non-carriers, implicating these as exposure-associated responses. IsdB-specific ADCD remained consistently high across all groups, underscoring its broad immunodominance. A key observation was the marked suppression of ADNP and ADCP in children with acute SSTI, with minimal phagocytic activity across all tested antigens. Although phagocytic activity partially recovered in SSTI at convalescence, ADNP and ADCP activity remained significantly lower than in convalescent invasive infection. However, during acute invasive disease, ADNP responses to SrtA and ClfAB and ADCP responses to Hla, HlgA, and SdrD/SdrE were all significantly reduced compared to carriers. Specific to the Hla, SdrD/SdrE, and HlgA ADCP response, convalescence from invasive disease was associated with high responses that mirror those observed in healthy children. These patterns suggest that despite repeated exposures, children with SSTI or acute invasive disease may fail to mount effective Fc-mediated clearance mechanisms, contributing to disease susceptibility.

### Multivariate modeling identifies polyfunctional antibody signatures that discriminate *S. aureus* clinical phenotypes

Having identified substantial differences in antibody levels, avidity, glycosylation and effector functions across distinct clinical states, we sought to define a minimal set of antibody features that reliably discriminates these phenotypes. To minimize overfitting, we applied a supervised machine-learning framework integrating least absolute shrinkage and selection operator (LASSO) feature selection with partial least squares discriminant analysis (PLS-DA). Comparison of healthy children harboring *S. aureus* with non-carriers revealed clear separation, with high cross-validation accuracy, driven by elevated ADCD to IsdB, LukAB, and SAK, and higher avidity and increased IgG1/IgG3 levels in colonized individuals. In contrast, healthy non-carriers were enriched for SpA-specific ADCD and higher avidity IgG responses and IgA1 to SdrE **(Figure S7)**. LASSO-selected features were further explored through correlation networks, which revealed structured clustering of a specific set of antigen-specific responses, including superantigens (SEA, SEB), EsxB, leukotoxins (Hla, HlgA, LukS), and surface proteins (IsdA, IsdB, SdrD), that collectively tracked with resilience to the carrier state. Finally, the violin plots show superior performance of the model with selected features over models with randomly selected features or permuted data labels, highlighting the robustness of the model to classify the target groups using the selected features.

The healthy carrier state exhibited a distinct *S.aureus* specific humoral immune profile when compared to children with invasive disease (**Figure 3a and b**). Carriers consistently exhibited higher ADCP and ADNP activity along with increased FcγR binding to conserved surface antigens (SdrD/SdrE, IsdA/IsdB, ClfB) and Hld. In contrast, SSTI cases were dominated by leukotoxin-focused responses. Correlation networks revealed antigen-specific clustering (e.g., ClfB, SdrD, SdrE), with distinct functional modules **(Figure 3a)**. These differences persisted when comparing healthy carriers with convalescent SSTI **(Figure 3b).**

**Figure 3.**
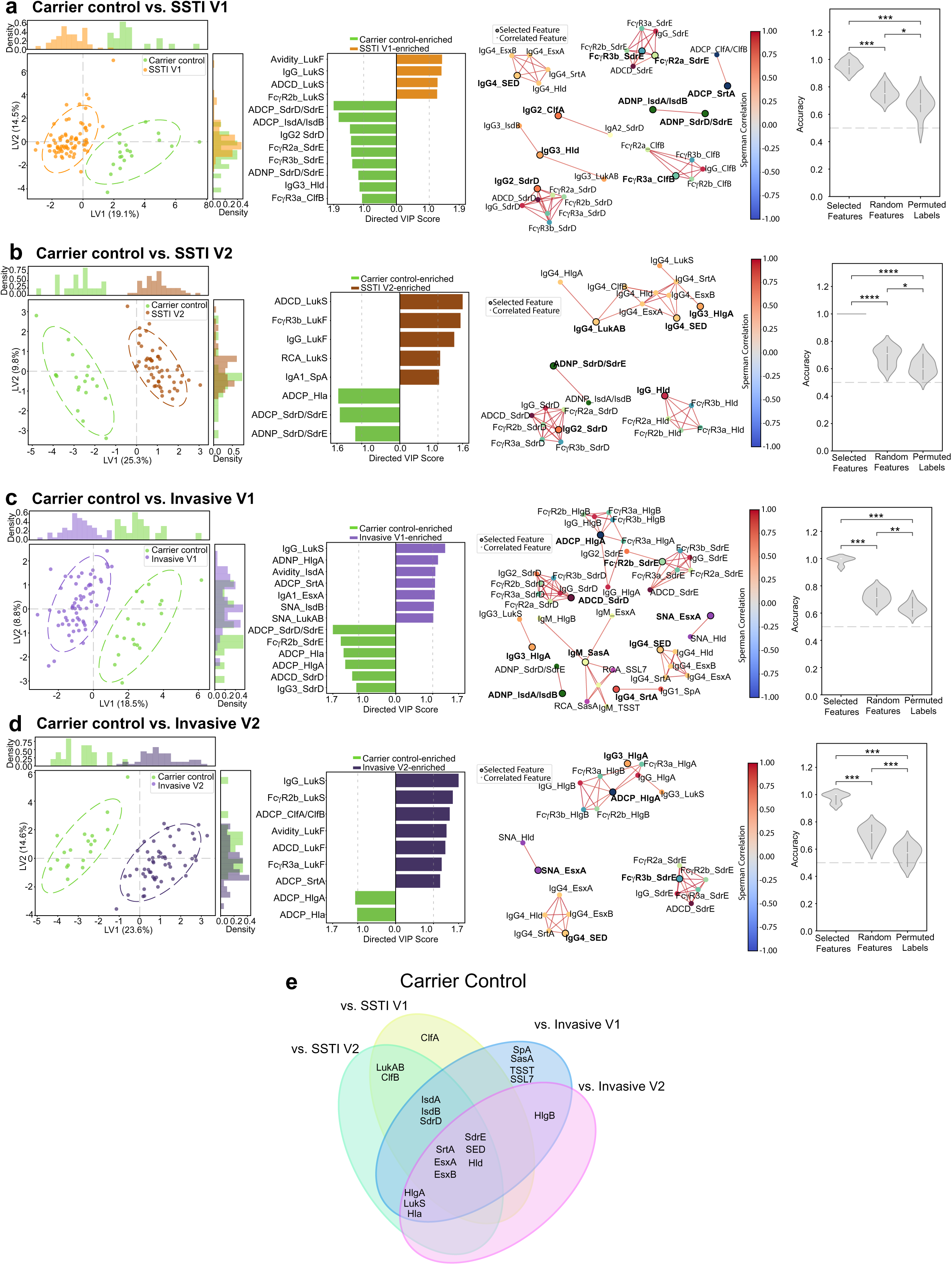
Multivariate modeling identifies minimal antibody features that distinguish carrier controls from disease. (a-d) Least absolute shrinkage and selection operator (LASSO) was used to identify minimal sets of antibody features that discriminate between carriers and various disease states. Multivariate modeling was performed using partial least squares discriminant analysis (PLS-DA) for each comparison: carrier control vs SSTI V1 (a), SSTI V2 (b), Invasive V1 (c), and Invasive V2 (d). For each comparison, plots from left to right show: PLS-DA score plots of the first two latent variables depicting sample separation; directional bar plots of variable importance in projection (VIP) scores for LASSO-selected features (VIP >1.0); correlation networks linking clinical groups to selected features (Spearman ρ > 0.8, FDR-adjusted p < 0.05); and violin plots showing model accuracy across 100 iterations of 5-fold cross-validation, benchmarked against models built from random features and permuted labels. (e) Venn diagram summarizing antibody features enriched in the carrier control across four pairwise comparisons (vs SSTI V1, SSTI V2, Invasive V1 and Invasive V2) based on LASSO selection and correlation network analysis.

Comparison with invasive disease yielded similar results – healthy carriers exhibited higher ADCP to Hla, HlgA, and SdrD/SdrE, enriched FcγR2b binding to SdrE, and elevated ADCD and IgG3 to SdrD **(Figure 3c)**. At convalescence from invasive disease, healthy carriers retained enhanced ADCP to Hla and HlgA **(Figure 3d)**. Cross-comparison identified a conserved set of features i.e. responses to SrtA, EsxA, EsxB, SdrE, SED, and Hld consistently enriched in healthy carriers, defining a polyfunctional signature potentially associated with protection against invasive disease **(Figure 3e)**. Moreover, cross-comparison illustrated critical differences between SSTI and invasive disease both in terms of the antigenic targets of the host response as well as their functional attributes. These findings suggest that exposure to *S. aureus* in the carrier state is associated with enriched functional antibody repertoires targeting conserved staphylococcal surface proteins and toxins, that are lost or not present upon skin or invasive disease. The dominance of inflammatory toxin-specific responses in disease may thus reflect immune activation secondary to the magnitude of antigen exposure and tissue damage rather than protective immunity per se.

### Antibody signatures associated with protection from recurrence

As 33% of children with SSTI and 8.6% of invasive cases in this cohort experienced recurrent *S. aureus* disease, we leveraged this opportunity to examine antibody features associated with protection. Recurrent SSTI cases displayed elevated IgG4 to IsdB, LukD, and SCIN, IgG2 to LukD, IgG to SrtA, IgM to SdrD and IsdB and FcγR2b binding capacity by SrtA-specific antibodies, collectively representing poorly functional humoral immune responses **(Figure 4a)**. In contrast, children without recurrent SSTI displayed broader, functionally optimized responses, including multi-isotypic antibodies to SEB, Hld, IsdA, IsdB and SdrE. These individuals also showed higher avidity ClfB antibodies and modestly increased SNA and RCA binding across most antigens, suggesting a shift toward anti-inflammatory glycoforms **(Figure 4b)**. Functionally, children without recurrent infection had higher ADCP activity against Hla, indicating that robust opsonophagocytic toxin clearance may be associated with protective immunity consistent with our prior studies^43, 44^ **(Figure 4c)**.

**Figure 4.**
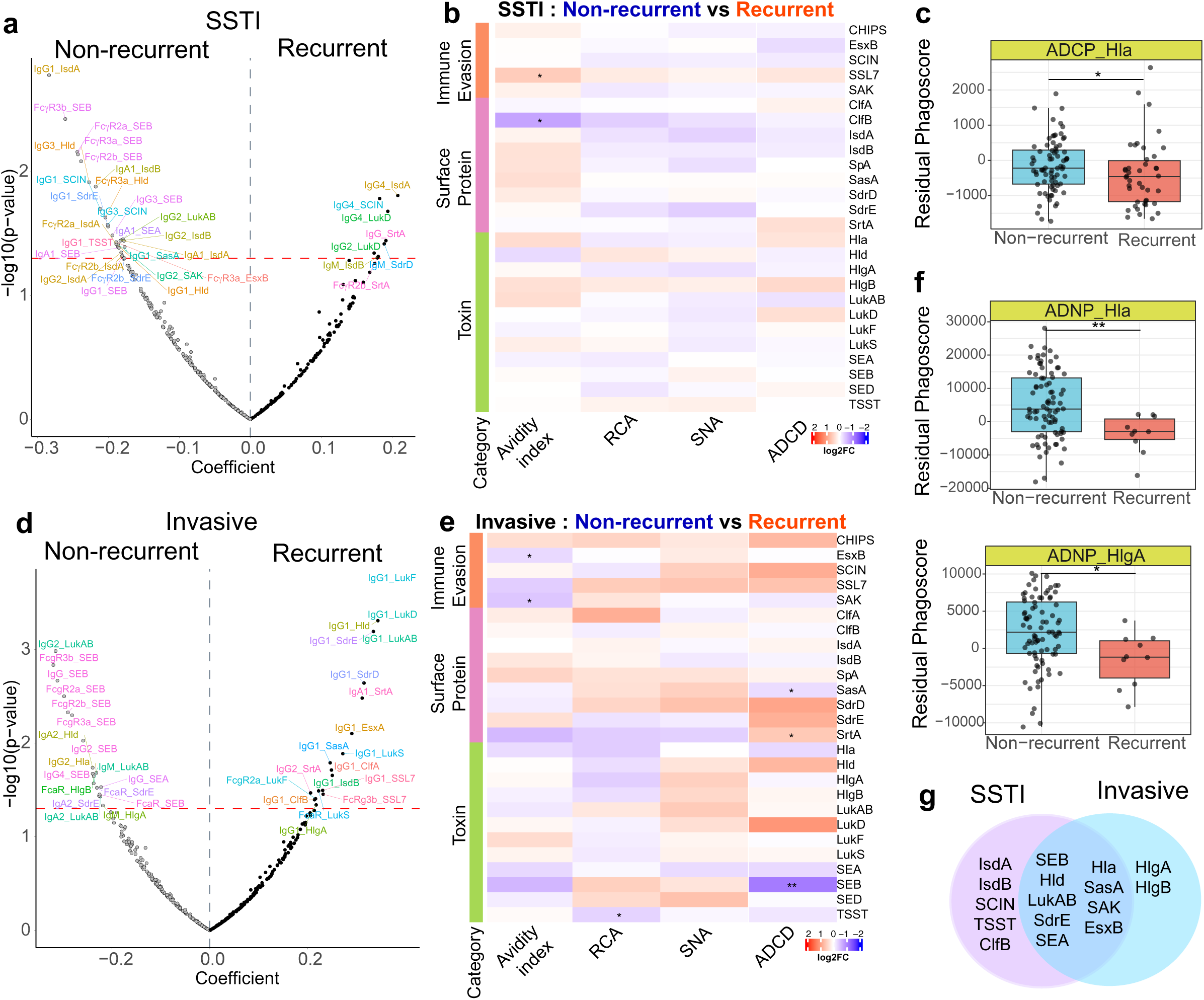
Distinct antibody profiles and functional responses are associated with recurrent SSTI and Invasive in pediatric *S. aureus* infections. (a,d) Volcano plots showing univariate comparisons of antigen-specific IgG1 and Fcγ receptor-binding features between non-recurrent and recurrent (a) SSTI or (d) Invasive patients after adjustment for age using residuals. Each dot represents an antibody feature, plotted by effect size (x-axis) and statistical significance (y-axis: –log10 p-value). Red dashed line denotes nominal p = 0.05. Features significantly enriched in the recurrent group are labeled and colored by antibody class and antigen specificity. (b,e) Heatmaps show the fold-change in antibody responses between non-recurrent and recurrent children within SSTI (b) or Invasive groups (e), calculated as the median difference in z-scored values for each antigen-feature pair. Rows represent *S. aureus* antigens grouped by functional class (immune evasion, surface proteins, and toxins), and columns represent antibody features including Avidity index, RCA binding, SNA binding and ADCD activity. Colors denote the direction and magnitude of fold change (red = higher in recurrent, blue = higher in non-recurrent), and stars indicate significance from Mann–Whitney U test (*p < 0.05, **p < 0.01, *p < 0.001). (c,f) Box plots showing residual phagocytic scores for ADCP and ADNP features enriched in non-recurrent SSTI (c) or Invaisve (f) individuals. Each point represents an individual; box plots show median and IQR. Residuals were derived from linear models adjusted for age. (g) Venn diagrams summarizing overlapping and unique antigens for which antibody features are enriched in non-recurrent SSTI and Invasive children.

In invasive disease, recurrence was associated with elevated IgG1 titers to multiple antigens, and increased FcαR and FcγR3b binding capacity by LukS and SSL7 antibodies, respectively. Additional features included higher IgG2 and IgA1 responses to SrtA. In contrast, children with invasive disease that did not manifest recurrence mounted broader, multi-isotypic responses including IgG2, IgA2, and IgM responses to LukAB, IgA2 and FcαR responses to SdrE, IgG to SEA, IgA2 to Hld, IgM to HlgA, and IgG2 to Hla. Protected children also demonstrated robust engagement of FcγRs and multiple isotype responses to SEB **(Figure 4d)**, higher-avidity IgG1 to EsxB and SAK, increased ADCD activity to SasA and SEB, and ADNP to Hla and HlgA **(Figure e-f)**. Shared analysis revealed protective antibody signatures targeting superantigens (SEA, SEB), leukotoxins (Hla, Hld, LukAB), immune evasion factors (SasA, SAK, EsxB), and surface proteins (SdrE) across both SSTI and invasive non-recurrent cohorts **(Figure 4g)**. These findings suggest that protection against recurrence is associated with broad, functionally active antibody responses targeting key bacterial virulence determinants.

### Hla neutralizing response serves as a biomarker of broad humoral immune breadth

Among serologic responses common to children without recurrent SSTI or invasive disease **(Figure 4g)**, the anti-Hla response is a known correlate of protective immunity among children^45^ and adults^46^. Prior work has shown higher maternal anti-Hla neutralizing antibodies (NAb) are associated with reduced infant risk of SSTI during the first year of life^19^, and experimental models demonstrate that Hla neutralization contributes to protection against multiple disease settings by preserving antigen-specific CD4^+^ T cell responses^32^ and enhancing neutrophil function^47^. However, whether these Hla-specific responses provide protection solely via neutralization or via additional functional mechanisms remained unknown, prompting us to selectively dissect the Hla-specific response. Hla IgG titers increased with disease severity, with children with invasive disease exhibiting significantly higher anti-Hla IgG titers and NAb titers compared with SSTI subjects and controls **(Figure 5a)**, mirroring patterns previously observed by luminex profiling **(Figure 1c)**. Anti-Hla IgG titers strongly correlated with neutralizing activity across individuals **(Figure 5b)**. Anti-Hla NAb titers were not significantly influenced by carrier status or recurrence **(Figure S8a-f)**; however children with convalescent invasive disease exhibited the highest NAb titers **(Figure S8d)**.

**Figure 5.**
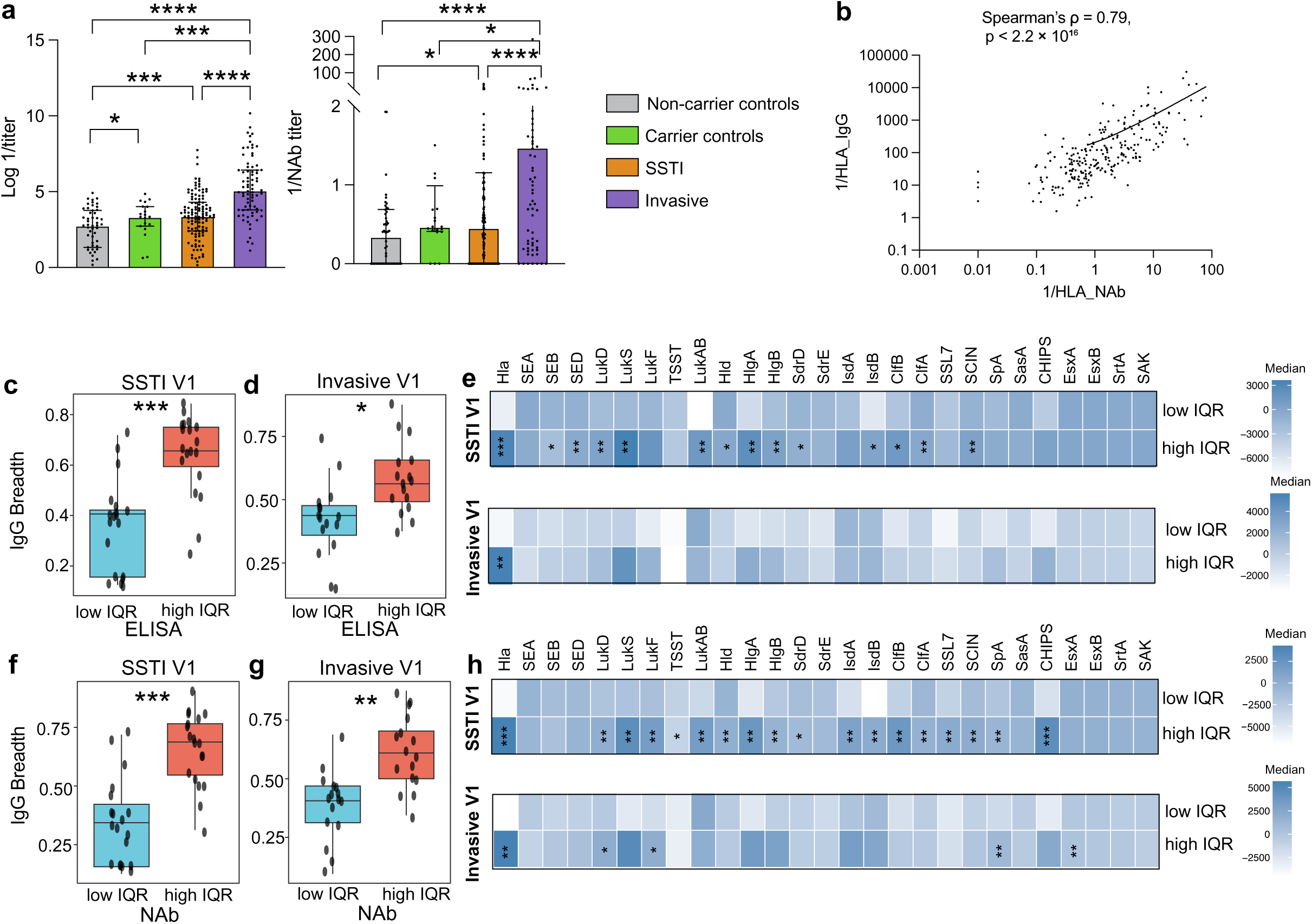
Anti-Hla IgG and NAb titers in children with SSTI and Invasive Infection. (a) Acute anti-Hla IgG measured by ELISA (left) and anti-Hla neutralizing antibody (NAb) titers (right) measured by RBC lysis assay in children with SSTI, invasive infection, carrier controls, and non-carrier controls. (b) Correlation between anti-Hla IgG and NAb titers across indviduals. Data expressed as log 1/OD450 or 1/NAb titer, comparisons by Wilcoxon rank-sum tests with Benjamini–Hochberg correction (**p < 0.05, **p < 0.01, ***p < 0.001). (c,d) Samples were stratified into low- and high-interquartile (IQR) groups based on anti-Hla ELISA titers. IgG breadth, defined as the fraction of antigen-specific responses exceeding the cohort median for each feature, was quantified using age-adjusted data in SSTI V1 (c) and Invasive V1 (d). (e) Heatmap shows age-adjusted median IgG responses to *S. aureus* antigens in SSTI V1 and Invasive V1 subjects stratified into low- and high-IQR groups based on anti-Hla ELISA titers. (f,g) Samples were stratified into low- and high-IQR groups based on anti-Hla NAb titers. IgG breadth was quantified using age-adjusted data in SSTI V1 (f) and Invasive V1 (g). (h) Heatmap shows age-adjusted median IgG responses to *S. aureus* antigens in SSTI V1 and Invasive V1 subjects stratified into low- and high-IQR groups based on anti-Hla ELISA titers. Boxplots indicate the median and interquartile range, with whiskers representing 1.5× IQR; points denote individual subjects. Statistical comparisons between groups were performed using Wilcoxon rank-sum tests with Benjamini–Hochberg correction (**p < 0.05, **p < 0.01, ***p < 0.001).

The protective activity of anti-toxin antibodies to anthrax is enhanced by Fc-mediated opsonophagocytic clearance^43, 44^. To examine whether Fc-biology could contribute to Hla-mediated protection, we next examined whether the magnitude of Hla-specific humoral response was associated with broader humoral immunity. We stratified children with SSTI and invasive infection into low- and high-Interquartile (IQR) groups based on anti-Hla IgG **(Figure 5c-e)** and NAb titers **(Figure 5f-h)**. High-IQR individuals, defined using either ELISA IgG or NAb titers exhibited significantly greater IgG breadth across *S. aureus* antigens in both SSTI and invasive cohorts **(Figure 5c-h)**. In addition, SSTI subjects in the high-IQR group displayed significantly higher breadth of FcγR-binding antibodies **(Figure S8g)**. While IQR stratification based on ELISA measurements did not reveal differences in IgG and FcγR-binding breadth at convalescence, Hla NAb-based stratification identified individuals with broader FcγR engagement, particularly among children with SSTI during convalescence **(Figure S8h-i)**. Together, these findings suggest that anti-Hla NAb titers may serve as a clinically-actionable biomarker to identify patients with broad anti-staphylococcal serologic responses. Protective anti-Hla NAb responses are observed in two distinct groups of infants/children: those whose naturally-occurring titer derived from maternal transplacental transfer or innocuous exposure affords protection against primary disease^19, 45^, and those who experience *S. aureus* infection, eliciting a robust and protective NAb response conferring resilience thereafter^45^. We hypothesize that elevated anti-Hla NAb titers in children may thus reflect not only protection from disease, but serve as a predictor of individuals with a coordinated, functionally broad humoral response to *S. aureus*.

### Correlate-guided Fc-engineering of IsdB antibodies results in enhanced bacterial control

To further determine whether additional functional humoral signatures underlie mechanistically useful correlates of immunity against *S.aureus,* we next focused on IsdB, for which imprinting had been observed in animal models resulting in poor bacterial protection^31^. These imprinted responses have been implicated in poor performance of the antigen in human vaccine studies^40, 41, 48^, thus here we sought to determine whether IsdB profiles associated with differential control of *S.aureus* in the PISA cohort could point to functional mechanisms to bolster IsdB-mediated protection against the bacteria. Thus, we generated a panel of Fc-engineered mAbs targeting IsdB. IsdB consistently ranked among the most enriched antigens in asymptomatic carriers and non-recurrent profiles and has demonstrated *in vivo* protection^13, 49^, despite failing as a vaccine antigen^50^. Whether this failure and reports of enhanced disease^51, 52^ reflects suboptimal Fc functionality remains unclear. Leveraging a validated IsdB-specific mAb^38^ to mimic functional profiles associated with asymptomatic infection, we engineered six Fc variants on a human IgG1 backbone to modulate FcγR interactions: wild-type (WT), Fc-silent (N297Q), and four Fc-enhanced forms (GASDALIE, SEHFST, EFTEA, and KWES).

All variants retained equivalent antigen-binding, confirming that Fc engineering did not alter IsdB recognition **(Figure 6a)**. FcγR binding profiles **(Figure 6b)**: N297Q showed minimal FcγR binding, GASDALIE exhibited broad and enhanced FcγRs binding. SEHFST selectively enhanced FcγR2a/2b engagement while KWES and EFTEA increased binding across FcγR2a, FcγR2b, and FcγR3a. Functional assays demonstrated that N297Q lacked activity across all effector assays, confirming the strict requirement for FcγR engagement **(Figure 6c-e)**. GASDALIE induced the strongest ADCP activity, consistent with its broad FcγR engagement, but exhibited reduced ADCD activity, suggesting a tradeoff between phagocytic and complement pathways. SEHFST enhanced both ADCP and ADCD, KWES preferentially augmented ADCD, and EFTEA elicited the highest ADNP and ADCD activity. Notably, EFTEA displayed the desirable profile, closely resembling that observed in carrier children and distinct from profiles in diseased individuals.

**Figure 6.**
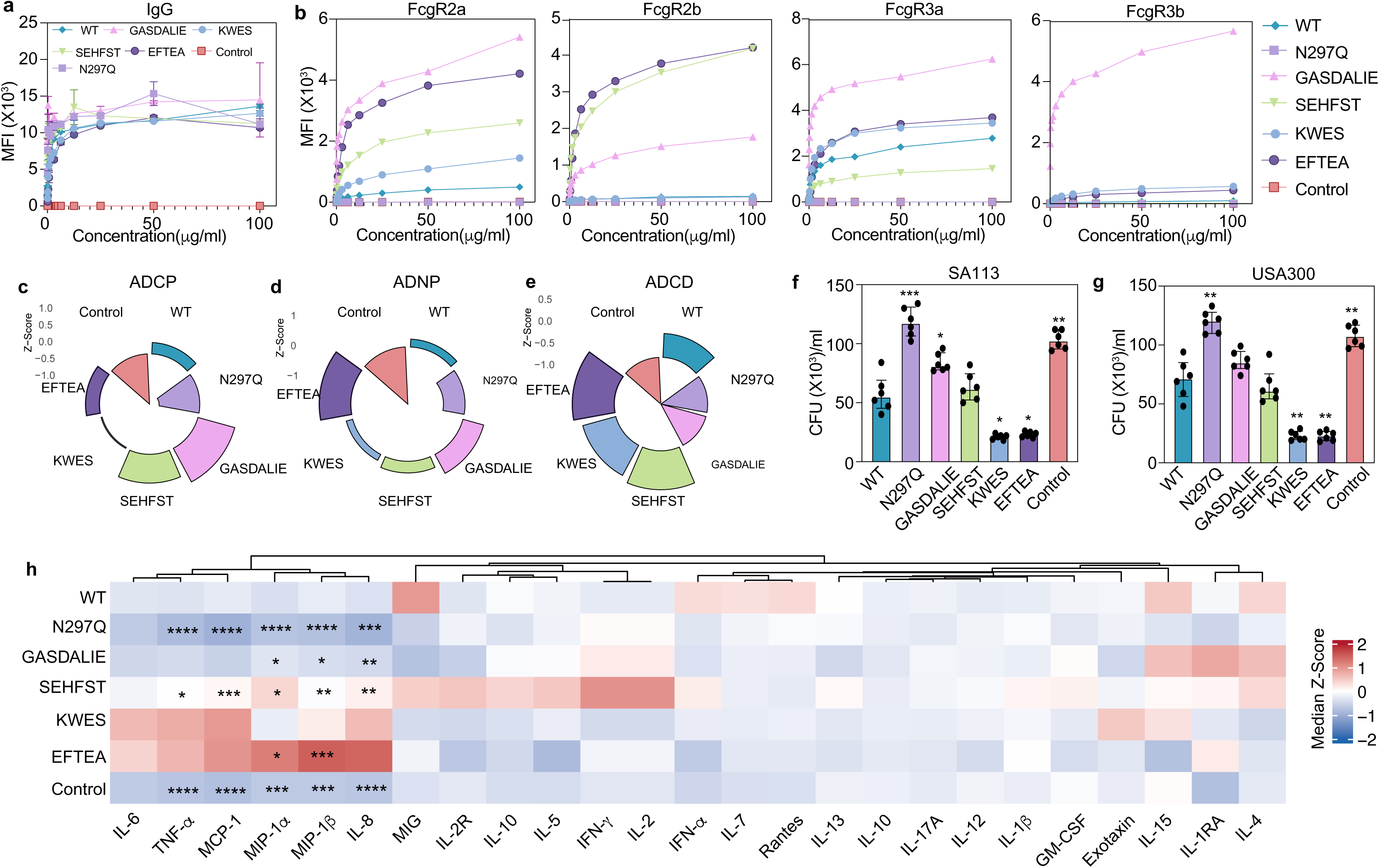
Functional characterization of Fc-engineered IsdB monoclonal antibody variants. (a) Binding kinetics of IsdB-specific mAb Fc variants to recombinant IsdB protein were measured by Luminex. Antibody variants included wild-type (WT), Fc-silent (N297Q), and Fc-enhanced (SEHFST, GASDALIE, EFTEA, and KWES) forms. Each point represents a replicate (n=2); lines connect mean ± SEM. (b) Binding kinetics of IsdB-specific mAb Fc variants to FcγR were measured by Luminex, lines connect mean (n=2). (c-e) Rose plots illustrating Z-scored effector functions of IsdB Fc variants across three effector functions: (c) ADCP, (d) ADNP, and (e) ADCD. Each segment represents one variant’s activity relative to the others. Control indicates irrevalant SARS-Cov2 mAb. (f-g) Opsonophagocytosis killing (OPK) assays were performed against *S. aureus strains* SA113 (f) and USA300 (g) using IsdB-specific mAbs bearing the indicated Fc modifications. Each dot represents an individual replicate; bars indicate median with interquartile range. Statistical significance was determined relative to WT antibody using one-way ANOVA with multiple comparisons (*p < 0.05, **p < 0.01, ***p < 0.001). (h) Heatmap showing median Z-scored cytokine and chemokine levels measured by Luminex in supernatants from OPK assay (g). Asterisks denote statistically significant differences relative to WT antibody calculated by one-way ANOVA with multiple comparisons (*P < 0.05, **P < 0.01, ***P < 0.001, ****P < 0.0001).

Opsonophagocytic killing (OPK) is regarded as the presumed correlate of immunity for surface-binding *S aureus* antibodies ^53^. Fc-variant testing in an OPK assay using the well-characterized SA113 and USA300 *S. aureus* clinical isolates revealed pronounced Fc-dependent differences in bacterial clearance, with KWES and EFTEA mediating the most effective bacterial killing **(Figure 6f-g)**. Furthermore, Luminex profiling of OPK supernatants further revealed increased levels of select pro-inflammatory and chemotactic mediators including IL-6, TNF-a, MCP-1, MIP-1a, MIP-1b, and IL-8 following treatment with KWES and EFTEA Fc-variants that may be necessary for T and NK cell recruitment required for bacterial control and clearance **(Figure 6h)**. Collectively, these findings demonstrate that Fc-engineering can selectively enhance specific antibody effector functions without compromising antigen binding. Moreover, Fc profiles enriched in healthy children, compared to those with disease, can be leveraged to improve bacterial clearance. The ability to precisely tune phagocytic and complement-mediated immunity provides a rational framework for next-generation toxin and/or surface specific *S.aureus* mAb development or to guide vaccine design against *S. aureus*, grounded in the protective mechanisms observed in naturally resilient children.

## Discussion

This study provides the first comprehensive functional antibody analysis against *S. aureus* in a pediatric cohort, offering critical insights into the immunological correlates of protection against disease and recurrent infection. Our findings extend current paradigms by showing that protection is determined not merely by antibody quantity, but by subclass distribution, avidity, Fc glycosylation, and effector function—parameters that shape functional immunity and disease outcome. The study also highlights actionable targets for rational development of risk-stratifying biomarkers as well as next-generation vaccine and antibody-based therapeutics.

Age-associated increases in anti-staphylococcal antibody responses reflect cumulative exposure and progressive immune maturation^54, 55^. A key finding of this study is the identification of qualitatively distinct antibody profiles of children that exhibit resilience against infection. Whereas SSTI and invasive disease were associated with broad, inflammatory, toxin-directed responses, colonization observed in the healthy carrier population correlated with focused, high-quality antibodies characterized by strong FcγR engagement, robust opsonophagocytosis, and complement deposition targeting conserved surface proteins (ClfB, IsdA/IsdB, SdrD/SdrE) and toxins (Hla, HlgA, LukAB). Enrichment of opsonic antibodies to SdrD and SdrE, adhesins that are critical for epithelial attachment and colonization^56^, suggests that blocking bacterial adherence may represent a key protective mechanism. Similarly, antibodies that interfere with the function of IsdB which is essential for heme-iron acquisition, and ClfB, which mediates adherence to fibrinogen and cytokeratin 10, may modulate colonization and therefore disease risk^14, 49, 57,13, 49, 58^. Antibodies against major toxins (Hla, HlgA, LukAB) likely confer additional benefit by neutralizing cytolytic activity thereby preserving the functional attributes of host innate and adaptive immune cells. Supporting this, immunization with detoxified Hla variants enhances T follicular helper responses in mice, linking toxin-neutralizing antibodies to improved adaptive immunity^59, 60^. Notably, each of these identified targets have shown protective efficacy in preclinical models^13, 14, 49, 61^, reinforcing their relevance as candidates for vaccine and therapeutic development.

The serologic responses observed in children who developed disease were characterized by increased antibody sialylation and galactosylation, glycoforms known to reduce FcγR binding and impair opsonophagocytic potential, as previously speculated due to imprinting^31^. At convalescence, antibody glycan profiles transitioned toward more inflammatory, effector-capable states, underscoring the principle of “Right antigen, Right Fc format.” These findings parallel murine data showing that *S. aureus* drives IL-10–mediated antibody sialylation, impairing neutrophil function and bias the antibody repertoire toward non-functional Fc profile^31, 62^. Mechanistically, our Fc-engineering experiments suggest only select Fc configurations supported robust opsonophagocytic killing, whereas other Fc variants, despite retaining antigen binding and opsonization capacity, failed to trigger effective microbicidal pathways. Such a mechanism could explain prior reports of enhanced disease associated with IsdB-targeting antibodies when Fc effector functions were improperly tuned^31, 51, 52^. The enrichment of low-sialylation antibodies in resilient children further implicates Fc glycan composition as a determinant of protection. Notably, while some antigens (ClfA, Hla, IsdB) exhibited hyper-sialylated antibodies during disease, others (ClfB, SdrD, SdrE) did not, indicating antigen-specific mechanisms of immune suppression.

Despite robust antibody titers following natural infection, ∼50% of patients remain susceptible to recurrence^63, 64, 65^ suggesting that antibody quantity alone does not predict protection. This conclusion is supported by failed *S. aureus* vaccine trials, all of which elicited high-level human antibody responses to each vaccine antigen evaluated. In our study, children with recurrent infection exhibited both quantitative and qualitative antibody defects, i.e. reduced avidity, and Fc engagement and effector functions to multiple key antigens. In contrast, children that did not experience recurrent infection mounted elevated levels of broad, multifunctional responses to key antigens (Hla, LukAB, SEA, SEB, SdrE, etc). The convergence of protective features in healthy children compared to SSTI and invasive disease suggests shared mechanisms of potential durable protective immunity. Inclusion of highly functional antibodies to toxins and superantigens (Hla, Hld, HlgA/B, LukAB, SEA, SEB, TSST) in the protective signature indicates that neutralization of immune-dysregulating toxins may be essential for long-term protection.

Our study illustrates the potential for serology-informed development of both preventive and therapeutic tools. Translation of the identified correlates of protection will require both the ability to identify individuals who are ‘at risk’ based on quantitative serologic stratification tools and the ability to develop active or passive immunization strategies that confer protection. This study provides early validation that these approaches are feasible. Building upon knowledge that the anti-Hla response is a correlate of protection against *S. aureus* disease^19, 45, 46^, we harness systems serology to reveal that those children with the highest anti-Hla titers also exhibit broad functional responses against multiple *S. aureus* antigens that contribute to resilience.

The identification of specific antigen-function combinations that correlate with natural protection provides a roadmap for rational vaccine and therapeutic design. Key principles include: (a) Multiantigen approaches: the diversity of protective targets identified suggests that successful vaccines and therapeutics will likely need to promote the generation of broad-based functional responses, achievable either by targeting multiple antigens spanning different functional classes (i.e., surface proteins and toxins) or by selective targeting of specific immunomodulatory virulence factors that preclude the host from developing a broad functional response (i.e., Hla and LukAB). (b) Disease state-specific targeting: key differences in the observed protective responses between healthy carriers and children with either SSTI or invasive disease indicate that protection against one clinical state may not necessarily equate with broad protection. (c) Functional optimization: Fc-engineering experiments demonstrate that antibody effector functions can be enhanced through targeted modifications. Similarly, as adjuvant strategies shape vaccine-elicited effector responses, preclinical studies should test whether specific formulations can be tailored to induce the protective Fc profile identified here.

In summary, this study demonstrates that pediatric immunity to *S. aureus* is governed by distinct qualitative and functional antibody traits. By systematically mapping these profiles across health and disease, we define potentially actionable correlates of protection that can inform rational vaccine and therapeutic design. Our focus on children provides critical insight on the heterogeneity of the human response to *S. aureus* in the context of both childhood development and early exposures to this microbe. However, it is important to recognize that the study of children is not likely to reflect the magnitude of immune imprinting that characterizes the host response to *S. aureus* in adults. Additionally, cellular immune responses and unmeasured antigens warrant further exploration, as the integrated function of the serologic and cell-based immune response may be key to bacterial control and prevention of disease. Collectively, these findings establish a comprehensive framework for understanding humoral protection and emphasize that antibody quality—rather than quantity—is a cornerstone of effective *S. aureus* immunity.

## METHODS

### Study Population, Sample Collection, and Longitudinal Follow-up

Pediatric participants were enrolled from St. Louis Children’s Hospital in St. Louis, Missouri, and Texas Children’s Hospital in Houston, TX in an ongoing prospective, observational cohort study beginning in August 2008 and June 2022, respectively.^45^ The study enrolled pediatric participants with the following characteristics: 1) healthy controls with or without asymptomatic *S. aureus* colonization, defined as carriers and non-carriers throughout the study, 2) children with culture-confirmed *S. aureus* SSTI, or 3) children hospitalized with an acute invasive *S. aureus* infection. Patients with traditional risk factors for healthcare-associated MRSA infections (e.g., indwelling catheter or percutaneous medical device, dialysis, post-operative infection, residence in a long-term care facility) were excluded from participation. Immunocompromised individuals were excluded, including preterm neonates, children with diabetes, renal disease, oncologic disease, or HIV. We excluded participants from whom *S. aureus* was recovered from a surface culture (e.g., decubitus ulcer) or for whom culture was not obtained. Participants who did not complete any follow-up visits were excluded from recurrent disease analysis. The Washington University and Baylor College of Medicine Institutional Review Boards approved study procedures. Informed consent, assent when appropriate, was obtained for all participants.

At enrollment, detailed questionnaires were administered to parents or guardians to document participant and family health history. For patients with active infection, medical charts were abstracted to obtain details regarding symptoms, culture results, laboratory and imaging results, and hospitalization outcomes. Colonization cultures (CultureSwab Liquid Stuart, Becton Dickinson, Franklin Lakes, NJ) were obtained from the anterior nares, axillae, and inguinal folds of each participant. *S. aureus* was identified from culture swabs by standard methods as previously described.^66^ An acute blood sample was drawn at enrollment from all participants and a convalescent blood sample was drawn 4-8 weeks after in participants with SSTI or invasive infection. A follow-up survey was administered at the convalescent visit to document the clinical course of the infection and to identify epidemiologic factors associated with *S. aureus* infection. All participants were followed longitudinally with surveys administered by mail, telephone, or e-mail every 3 months for 1 year to document interval infections and antimicrobial use. A recurrent infection was defined as a clinically diagnosed abscess, boil, cellulitis, or *S. aureus* culture-confirmed invasive infection (e.g., bacteremia, bone or joint infection). Participant information was collected and managed in REDCap electronic data capture tools hosted at Washington University in St. Louis.^67, 68^ Clinical Metadata were recorded **(Table S1)**. These samples were deidentified at Washington University and then shipped to the Ragon Institute of Massachusetts General Hospital, Massachusetts Institute of Technology and Harvard. Samples remained blinded to individuals conducting experiments until all data had been collected, quality-controlled, and integrated.

### Luminex-Based Systems Serology

Custom Luminex MagPlex bead-based multiplex assays were used to quantify antibody isotypes and subclasses (IgG, IgG1-4, IgA1, IgA2 and IgM), and Fc receptor (FcγR2a, FcγR2b, FcγR3a, FcγR3b, FcαR) binding across 27 canonical *S. aureus* antigens and 5 control antigens (Influenza HA, EBOV GP, Diphtheria Toxoid, Hepatitis B, Albumin). An equal mixture of influenza antigens from HA1 from different strains (Immune Technology Corp) was used as a positive control, and ebolavirus glycoprotein (Sino Biologicals) was used as a negative control. Additional controls included Diptheria toxin (Native antigen), Hepatitis B Surface Antigen (HepB, Creative Biomart) and Albumin (Sigma). Staphylokinase (SAK), EsxA, EsxB, SCIN, SSL7, CHIPS, IsdA, SdrD, SasA, Sortase A (SrtA), Protein A (SpA) and Hla were provided by the Bubeck Wardenburg laboratory. For SpA, SpA_KKAA_ mutant was used and for Hla, the Hla_H35L_ mutant protein was used^9, 69^. Staphylococcal Enterotoxin D (SED) and SdrE (53-601aa) was purchased from Cusabio. Panton-Valentine Leukocidin LukS (LukS), Panton-Valentine Leukocidin LukF (LukF), LukAB, LukD, HlgA, HlgB, TSST-1, Hld, SdrE and ClfA were purchased from IBT bioservices. ClfB, Staphylococcal Enterotoxin A (SEA), Staphylococcal Enterotoxin B (SEB) were purchased from MEDCHEM EXPRESS **(Table S3)**.

Carboxylated magnetic beads (Luminex Corp.) were covalently coupled to each antigen using standard EDC–Sulfo-NHS (Thermo Fisher Scientific) chemistry as per the manufacturer’s instructions. Samples were heat inactiavated at 56°C for 30 min and assays were optimized over a dilution curve to ensure selection of a dilution within the linear range of the assays. Diluted serum samples (1:30 for IgG and IgG1, 1:8 for IgG2 and IgG4; 1:5 for IgG3, 1:20 for IgG3, 1:10 for IgA1, IgA2, 1:5 for IgM; 1:15 for FcγRs; 1:10 for FcαR) were incubated with antigen-coupled beads overnight at 4°C. For subclass and isotype detection, PE-conjugated secondary antibodies (Sourthern Biotech) were used. Fc receptor binding was assessed using biotinylated recombinant FcγRs (DukeUniversity Protein Production Core) using BirA (Avidity) and conjugated to streptavidin-PE (PhycoLink). All secondary incubations were performed over 1 hour at room temperature. For measurements of avidity index, post-immune complex formation, samples were incubated with PBS with 0.1% BSA and 0.05% Tween with or without 7 M urea for 20 min and then washed three times before incubation with secondary detection reagents. Avidity index was calculated as the ratio of MFI in the presence ofurea treatment to that without urea treatment^70^. To assess antibody sialytion and galactosylatin, coupled antigens were incubated with 1:3 diluted serum overnight at 4°C to allow immune complex formation, followed by addition of fluorescein-labeled plant based lectin detects, SNA (1:100) and RCAI (1:500) (Vendor Laboratories) for 1 hour at room temperature. Median fluorescence intensity (MFI) (Prozyme) for each bead region was acquired using a xMAP Intelliflex (Luminex Corp). All samples were assayed in duplicate, and values were averaged followed by substraction of PBS background signal.

### Quantification and statistical analysis

All assays were run in duplicate and measurements were averaged for each sample and PBS substracted. All Luminex data were z-scored across samples for visualization and multivariate analysis. Group comparisons were performed using Mann-Whitney U tests. Age-adjusted univariate analyses were conducted using linear regression with age as a covariate. Volcano plots show effect size (regression coefficient) vs. statistical significance (−log₁₀p-value), with p < 0.05 as the nominal threshold. Correlation analyses used Pearson’s R. All analyses were conducted in R (Version 2024.12.1+563).

## Acknowledgments

We thank T. Ragon and S. Ragon and the Ragon Institute of MGH, MIT and Harvard for support. We thank Andrea Forbes, RN, and Linda Lamberth for assistance with participant recruitment and sample processing.

## Author contributions

M.Z.K, G.A., B.J. and J.B.W conceptualized the study. Luminex-based measurements of isotype and subclass concentrations, FcR binding, avidity, and all functional assays were performed by M.Z.K and T.S. OPK experiment was performed by M.Z.K. C.K. performed anti-Hla antibody measurement by ELISA and RBC lysis assay. M.Z.K, W.J. and E.K. performed all analysis and created all data visualizations under the supervision of B.J, D.L, and G.A. Samples were collected and metadata was curated by C.K, S.A.F, J.M., K.H., S.K., and J.B.W., M.G.B, L.F, J.P, J.N.W., J.C.M, K.G.H., and S.L.K. contributed to collection of human samples and data management. Project administration was performed by L.F and M.G.B. The manuscript was drafted by M.Z.K, C.K., G.A., B.J., and J.B.W. and reviewed and edited by all authors.

## Data and materials availability

Metadata associated with this study is provided in Table S4.

---

**Table S1.** Data dictionary defining clinical variables and coding schema used in PISA cohort.

| Variable | Description | Coding |
| --- | --- | --- |
| PPID | Study subject ID |  |
| Infxntype | Type of infection or control | 0, Healthy non-carrier \| 1, Healthy carrier \| 2, SSTI \| 3, Invasive infection |
| Age | Years |  |
| Sex | Sex | 1, Male \| 2, Female |
| Race | Race | 1, White \| 2, Black \| 3, Asian \| 4, Pacific Islander \| 5, American Indian/Alaskan Native \| 6, Other \| 7, Prefer Not to Answer/Unknown |
| Ethnicity | Ethnicity Hispanic | 0, No \| 1, Yes |
| IndexPriorStaph | Has your child ever been diagnosed with a *S. aureus* infection? | 0, No \| 1, Yes \| 2, Don't know |
| Index_infectionstrain | What type of staph infection did the index pt have at enrollment (SSTI and invasive groups only) | 1, MRSA \| 2, MSSA |
| Colonization_enrollment | Positive skin swab for *S. aureus* from any site | 0, No \| 1, Yes |
| Recurrence | Any recurrent SSTI during longitudinal follow-up? | 0, No \| 1, Yes |

**Table S2.** Participant Characteristics.

|  | Non-Colonized Control (n=58) | Colonized Control (n=20) | SSTI (n=152) | Invasive Infection (n=89) | P-value<sup>a</sup> |
| --- | --- | --- | --- | --- | --- |
| **Age** (mean, SD) | 2.6 (1.7) | 2.8 (1.8) | 2.4 (2.1) | 6.3 (5.3) | **p<0.001** |
| **Sex** |  |  |  |  | **p=0.004** |
| Male | 32 (55%) | 11 (55%) | 59 (39%) | 55 (62%) |  |
| **Race** |  |  |  |  |  |
| White | 51 (88%) | 16 (80%) | 89 (59%) | 48 (54%) | **p=0.014** |
| Black | 7 (12%) | 4 (20%) | 53 (35%) | 37 (42%) |  |
| Asian/Pacific Islander | 0 (0%) | 0 (0%) | 3 (2%) | 2 (2%) |  |
| Other | 0 (0%) | 0 (0%) | 6 (4%) | 2 (2%) |  |
| Prefer not to answer | 0 (0%) | 0 (0%) | 1 (<1%) | 0 (0%) |  |
| **Ethnicity** |  |  |  |  |  |
| Hispanic | 3 (5%) | 2 (10%) | 16 (10%) | 6 (7%) | p=0.76 |
| **Colonization at enrollment** |  |  |  |  |  |
| Yes | 0 (0%) | 20 (100%) | 92 (61%) | 44 (49%) | **p<0.001** |
| Unknown | 0 (0%) | 0 (0%) | 0 (0%) | 2 (2%) |  |
| **Infecting organism** |  |  |  |  |  |
| MRSA | N/A | N/A | 110 (72%) | 37 (42%) | **p<0.001** |
| MSSA |  |  | 41 (27%) | 50 (56%) |  |
| MRSA/MSSA |  |  | 0 (0%) | 1 (1%) |  |
| Unknown |  |  | 1 (1%) | 1 (1%) |  |
| **History of prior *S. aureus* infection** | N/A | N/A | 57 (38%) | 15 (17%) | **p<0.001** |

<sup>a</sup>Chi-square or one-way ANOVA comparisons between columns

**Table S1. Data dictionary defining clinical variables and coding schema used in PISA cohort.**

**Table S2. Patients Characteristics**

**Table S3. Antigen panel used for *S. aureus* systems serology profiling**

**Table S4. Systems Serology Metadata**

## Supplemental Methods

### Antibody-Dependent Cellular Phagocytosis (ADCP) and Antibody-Dependent Neutrophil Phagocytosis (ADNP)

Bead-based assays were used to quantify ADCP and ADNP, as previously described^1^. SdrE and SdrD; IsdA and IsdB; ClfA and ClfB were mixed in equal concentration for each family class. SdrD/SdrE, IsdA/IsdB, ClfA/ClfB, SrtA, HlgA and HLA were biotinylated using Sulfo-NHS-LC-LC biotin (Thermo Fisher, Waltham, MA, USA). A set of 2 antigens were mixed for multiplexing-SdrD/SdrE and IsdA/IsdB were coupled to custom-made Scarlett and Red fluorescent neutravidin beads (Thermo Fisher). Similarly, ClfA/ClfB and SrtA were coupled to custom-made Scarlett and Red fluorescent neutravidin beads, respectively and HlgA and HLA were coupled to custom-made Scarlett and Red fluorescent neutravidin beads, respectively. The bead mix and incubated with diluted plasma (1:80) to allow immune complex formation for 2h at 37°C. To assess the ability of sample antibodies to induce monocyte phagocytosis, THP-1s (ATCC) were added to the immune complexes at 1.25E5cells/ml and incubated for 16 h at 37°C. For ADNP, primary neutrophils were isolated by ACK lysis (Quality Biological) from whole blood. Isolated neutrophils at a concentration of 50,000 per well were incubated with immune complexes for 1h incubation at 37°C. Neutrophils were stained with an anti-CD66b PacBlue detection antibody (Biolegend) and fixed with 4% paraformaldehyde (Santa Cruz). Assays were acquired via flow cytometry with iQue (Intellicyt) and an S-Lab 384-well plate handling robot (PAA). For ADCP, events were gated on singlets and bead-positive cells. For ADNP, neutrophils were defined as CD66b positive events followed by gating on bead-positive neutrophils. A phagocytosis score was calculated for ADCP and ADNP as (percentage of bead-positive cells) x (MFI of bead-positive cells) divided by 10,000. Each sample was assayed in two independent technical replicates and averaged after subtraction of background signal from PBS controls.

### Antibody-Dependent Complement Deposition (ADCD)

To assess complement activation, antigen-coupled Luminex beads were incubated with serum (1:10) for 1 hour at 37°C, followed by guinea pig complement (Cedarlane) diluted 1:50 in gelatin veronal buffer (Boston BioProducts). After 30 minutes at 37°C, deposition of C3 was detected using a PE-conjugated anti-guinea pig C3 antibody. Data were acquired by Luminex and reported as MFI of C3 signal per antigen. MFI from technical duplicates were averaged after subtraction of background signal from PBS controls.

### IsdB Monoclonal Antibody and Fc Variants

The sequence of the IsdB-specific monoclonal antibody (mAb) was obtained from a previously published study^2^. Wild-type IsdB mAb and Fc-engineered variants were synthesized and purified commercially (Genscript). Antigen binding and Fc receptor engagement were quantified using a custom Luminex-based binding assay. Briefly, IsdB-coupled microspheres were incubated with serial dilutions of each mAb variant, and binding was detected with a PE-conjugated anti-human IgG or FcgRs, as discussed previously^3^. Functional activity of the IsdB mAb and its Fc variants was assessed by ADCP using THP-1 cells, ADNP using primary neutrophils, and ADCD using guinea pig complement^3^.

### Antigen Breadth Analysis

Antigen breadth was calculated to quantify the diversity of antigen-specific IgG1 responses across study groups. Normalized Luminex data were log-transformed and reorganized into long format, separating assay type and antigen identity. Median response values for each antigen were used as the threshold for positivity. For each sample, antigen breadth was defined as the proportion of antigens with signal intensity above the median value across all tested antigens within the same assay class. Breadth scores were computed for each individual and compared between clinical groups using nonparametric statistical tests.

### Feature Selection via Stability LASSO

Prior to analysis, all antibody feature data were log-transformed (if not already) and Z-score normalized using scikit-learn’s StandardScaler. To avoid overfitting and identify robust features, we implemented a repeated, stability-based Least Absolute Shrinkage and Selection Operator (LASSO) approach. To identify a minimal set of features that best explain group differences without overfitting, a five-fold cross-validation framework was implemented. The dataset was partitioned into five equally sized folds. For each repetition of the validation process, four folds (80% of the data) were used as a training set to build the model, while the remaining holdout fold (20% of the data) was used as a test set to evaluate its predictive performance. The performance for each model was measured by its classification accuracy on the unseen test data. This process was repeated five times, with each fold used as the test set exactly once, ensuring a robust and unbiased estimate of the model’s generalizability. The stability of each feature’s selection was then assessed; only features selected in over 80% of the iterations were retained for the final model.

### Classification and Visualization with PLS-DA

The features selected by the stability LASSO were used to build a Partial Least Squares Discriminant Analysis (PLS-DA) model. The model was constructed using the PLSRegression module from scikit-learn, regressing the selected features against the binary class labels. Model performance was assessed through 5-fold stratified cross-validation, with receiver operating characteristic (ROC) curves generated using sklearn.metrics.roc_curve. The performance of the final PLS-DA model was evaluated against two negative controls using a permutation testing framework: 1) a model built using an equal number of randomly selected features, and 2) a model built using the selected features but with permuted class labels. Statistical significance between the accuracies of the true model and the control models was assessed using the Mann-Whitney U test (scipy.stats.mannwhitneyu).

The contribution of each selected feature to the model was quantified using Variable Importance in Projection (VIP) scores. Features with a VIP score > 1.0 were considered particularly influential for group separation. To visualize the directionality of these associations, a directional VIP bar plot was generated. The sample-wise scores from the first two LVs of the PLS-DA model were plotted to visualize group separation, with 95% confidence ellipses to represent the distribution of each group.

### Correlation Network Analysis

To explain the co-regulation and interconnectedness of the humoral immune response beyond the LASSO-selected features, we performed a correlation network analysis. For each clinical group separately, we calculated all pairwise Spearman correlation coefficients between the LASSO-selected features enriched in that group and all other measured features.

The resulting p-values were adjusted for multiple comparisons using the Benjamini-Hochberg false discovery rate (FDR) correction (fdr = 0.05). A network was constructed where nodes represent immune features. Nodes were colored based on their antibody isotype or function. An edge was drawn between two nodes only if their correlation was statistically significant (FDR-adjusted p-value < 0.05) and the absolute correlation coefficient exceeded a threshold of |r| > 0.7. Two network visualization styles were used: a circular layout for a global overview and a force-directed (spring) layout to highlight tightly interconnected clusters of features. Network analysis and visualization were performed using networkx and matplotlib libraries in Python.

### Anti-Hla Neutralization Antibody Assay

Anti-Hla neutralization was determined as previously described{Tomaszewski, 2024 #301. Briefly, sera is diluted starting at 1:2 and incubated for 15 minutes with 2nM purified Hla at room temperature. 5×10^7^ washed rabbit red blood cells are added for 1 hour at room temperature. All specimens were tested in duplicate and in the presence of erythrocytes and without erythrocytes as background control. Cells and debris are centrifuged at 2000rpm for 10 minutes and supernatant measured for hemolysis (450nm, Tecan Infinite M200 Pro). A reference standard curve was generated with two-fold serial dilutions of a humanized monoclonal antibody against Hla (7B8) for each run (4-6 plates/run).^4^ 7B8 mAb was generated utilizing hybridomas from mice immunized with Hla_H35L_, a variant toxin with a single amino acid substitution which interferes with formation of the cytolytic pore.^5^ The log of the 7B8 standard reciprocal dilution is graphed against specific lysis readouts and the EC50 titer defined as the reciprocal dilution which neutralizes the toxicity of Hla by 50% using non-linear regression curves with log(inhibitory) vs. response-variable slope curves. Individual patient values are extrapolated to determine the Nab titer dilution that correlates with 50% lysis by the standard and the reciprocal value graphed. Background values are subtracted from each sample value prior to analysis. An assay negative control PBS-only well and positive control Triton X-100 only well are run with each set of plates.

### Anti-Hla Binding Antibody Assay

High-binding 384-well plates (Fisher Scientific) are coated with purified Hla at 1µg/mL in PBS overnight at 4°C. Plates are blocked with 0.1% BSA in PBS for 1 hour at room temperature. Serum sample dilutions added starting at 1:2 dilution and then three-fold for 24 dilutions in duplicate for 2 hours at room temperature. A standard curve using 7B8 Hla mAb was utilized and individual values extrapolated from a standard curve. All specimens were tested in duplicate. Plates are washed three times with 0.05% Tween 20/PBS prior to the addition of HRP-conjugated goat anti-human antibody (Southern Biotech) at 1:20,000 dilution for 45 min at room temperature. Plates are then washed five times, developed with TMB substrate (ThermoFisher) for 15 minutes and reactions are stopped with H_2_SO_4_. Absorbance (OD_450_) is measured on a Tecan plate reader.

---

**Figure S1.**
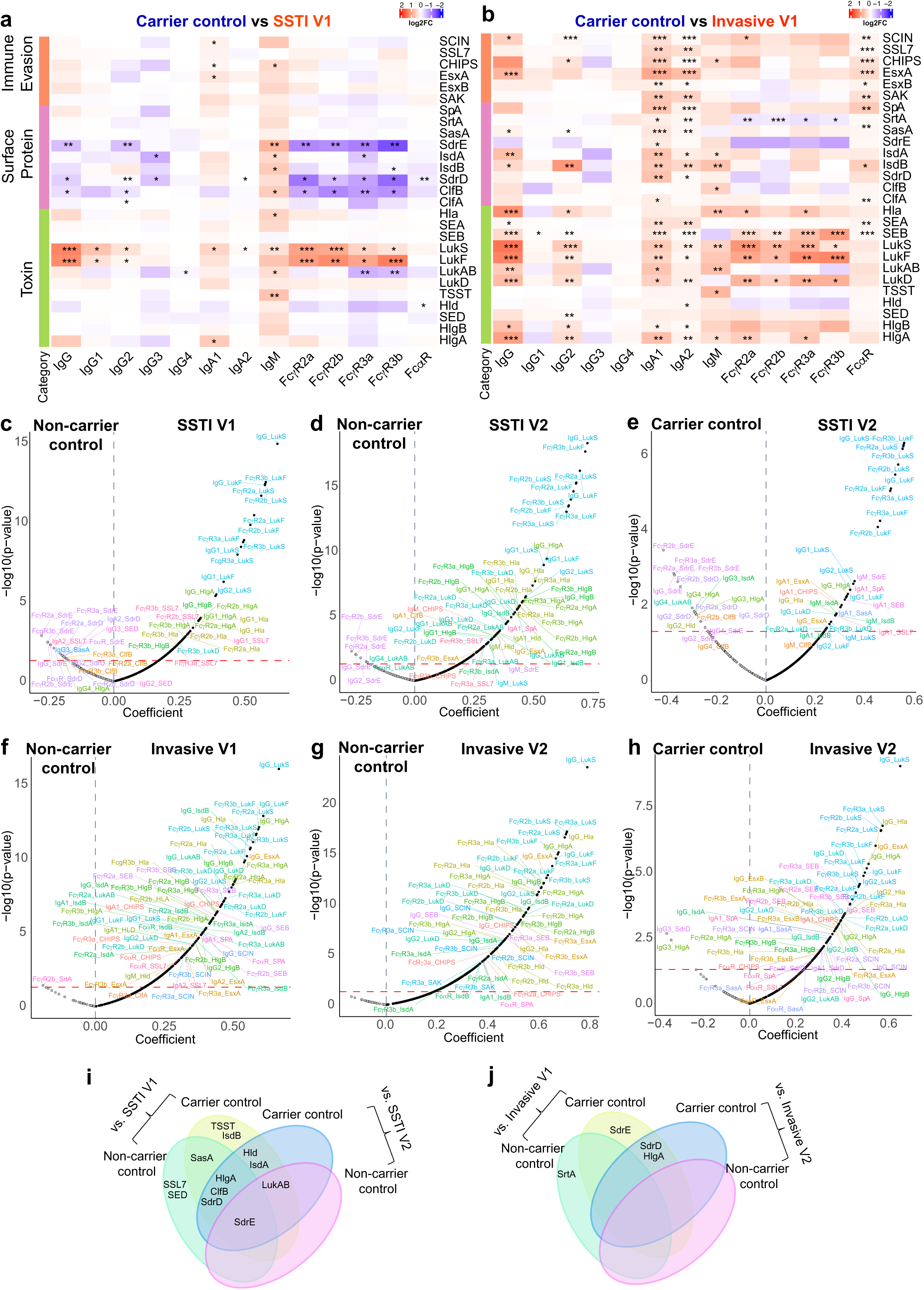
Distinct Antibody Features Discriminating Colonization from SSTI and Invasive Disease. (a-b) Heatmaps show the fold-change in antibody responses between healthy carrier children and those with acute SSTI (a) or acute invasive infection (b), calculated as the median difference in z-scored values for each antigen-feature pair. Rows represent *S. aureus* antigens grouped by functional class (immune evasion, surface proteins, and toxins), and columns represent antibody features including isotypes, subclasses, and Fc receptor binding profiles. Colors denote the direction and magnitude of fold change (red = higher in disease, blue = higher in colonized), and stars indicate significance from Mann-Whitney U test (*p < 0.05, **p < 0.01, *p < 0.001). (c-h) Volcano plots showing univariate comparisons between groups. Each point represents an antibody feature plotted by effect size (x-axis: regression coefficient) and significance (y-axis: –log10 p-value). Features significantly enriched in non-carrier control vs. SSTI V1 (c), non-carrier control vs. SSTI V2 (d), carrier control vs. SSTI V2 disease (e), non-carrier control vs. Invasive V1 (f), non-carrier control vs. Invasive V2 (g), carrier control vs. Invasive V2 disease (h) are labeled. The horizontal red dashed line marks significance (p = 0.05). Features are colored by antigen specificity. (i-j) 4-way Venn diagram summarizing features significantly enriched in non-carrier and carrier control individuals compared to SSTI (i) or Invasive (j).

**Figure S2.**
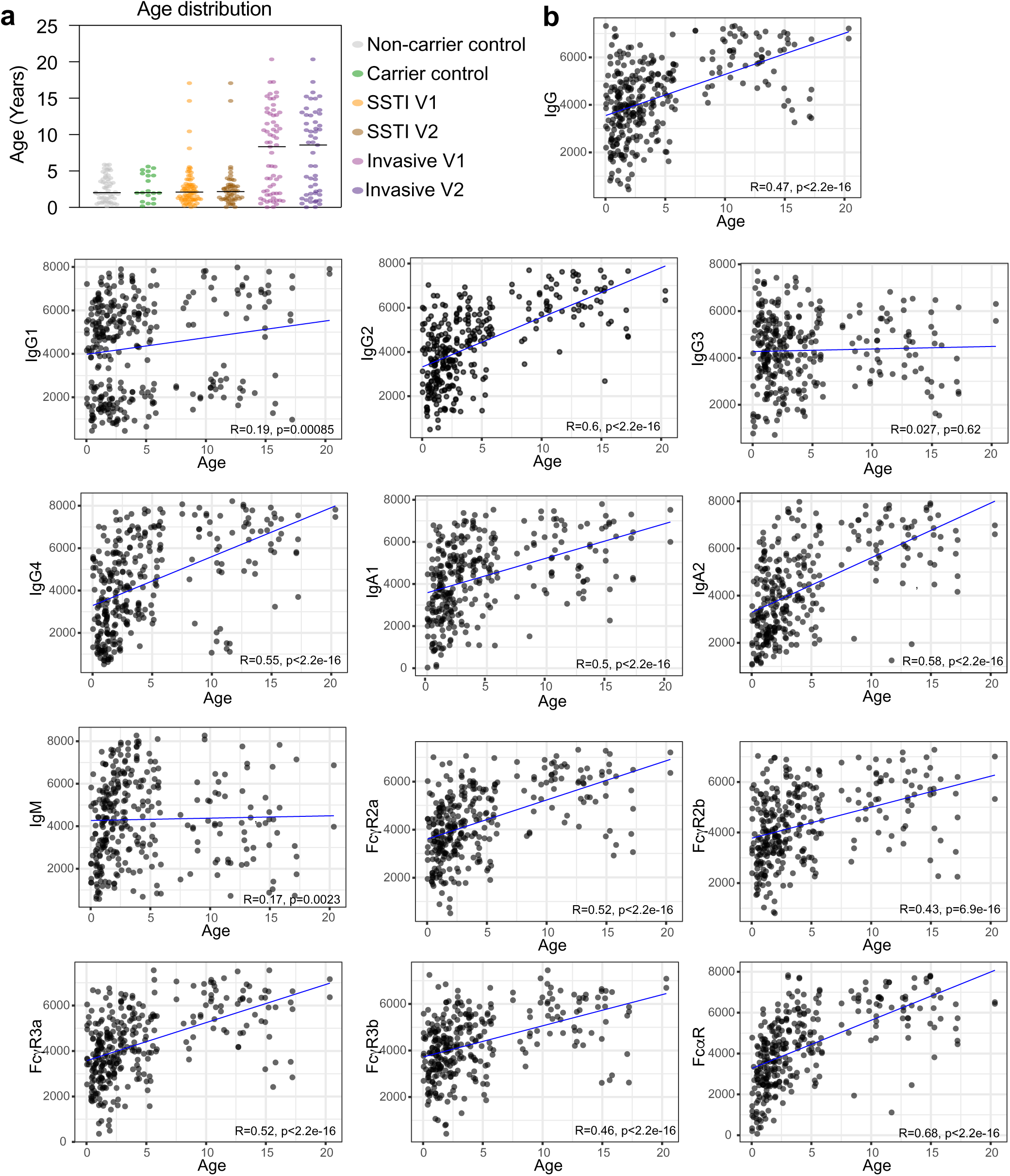
Age-Dependent Maturation of Humoral Immunity Across Pediatric *S. aureus* Infections. (a) Age distribution of subjects across study groups (healthy non-carrier and carrier control, SSTI V1/V2, Invasive V1/V2). (b) Scatterplots depict the relationship between age (x-axis, years) and total antibody levels or Fc receptor binding (y-axis, MFI) across all subjects. Each dot represents a subject. Spearman correlation coefficients (R) and *p*-values are shown.

**Figure S3.**
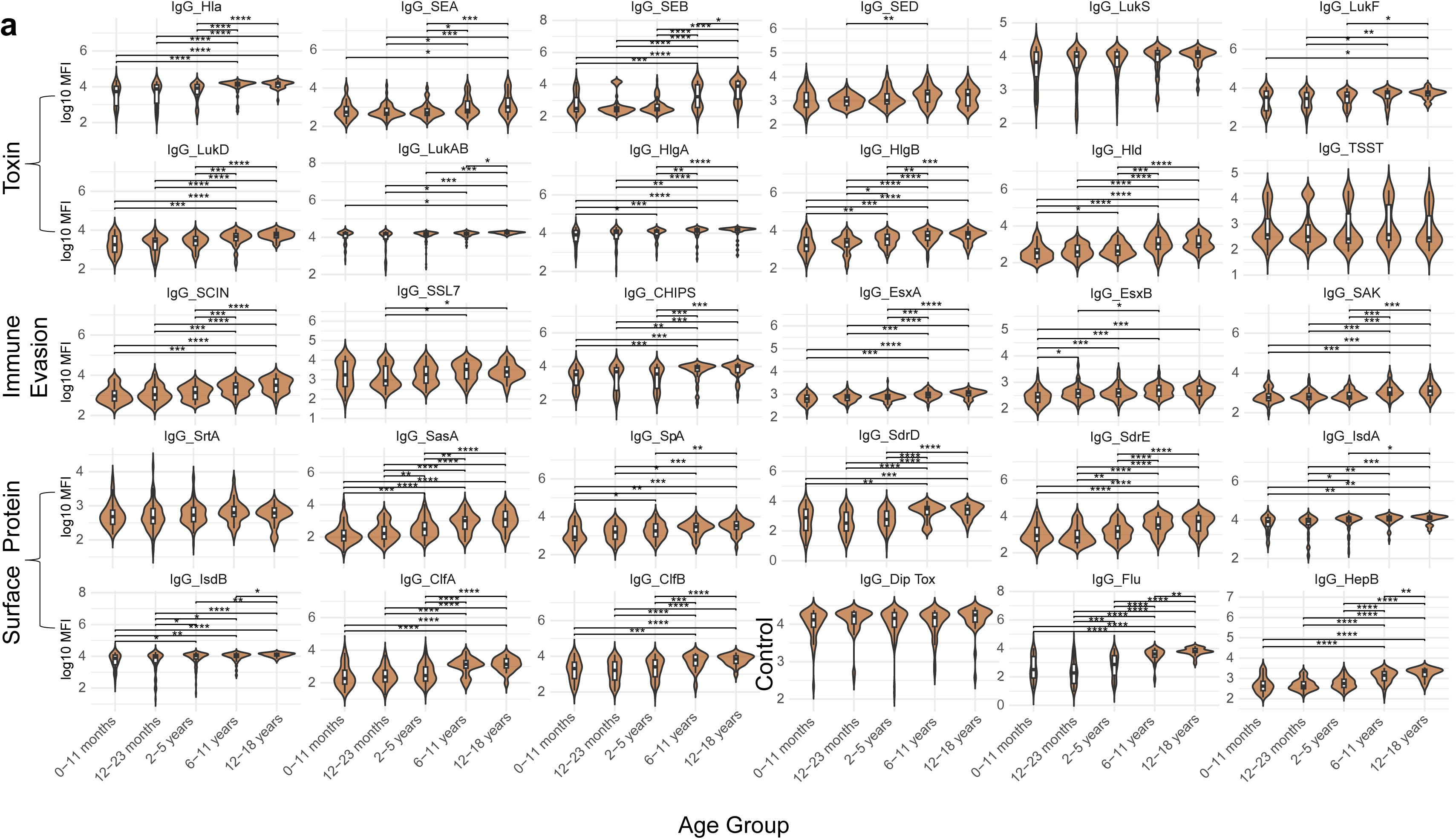
Age-Associated Increases in Antigen-Specific IgG Responses Across Development. (a) Violin plots with overlaid boxplots show log-transformed IgG titers to *S. aureus* antigens across five age-defined groups: 0–11 months, 12–23 months, 2–5 years, 6–11 years, and 12–18 years. Statistical significance for pairwise comparisons was assessed using the Mann-Whitney U test, with p-values denoted as follows: p < 0.05 (*), p < 0.01 (**), p < 0.001 (***), p<0.0001 (****).

**Figure S4.**
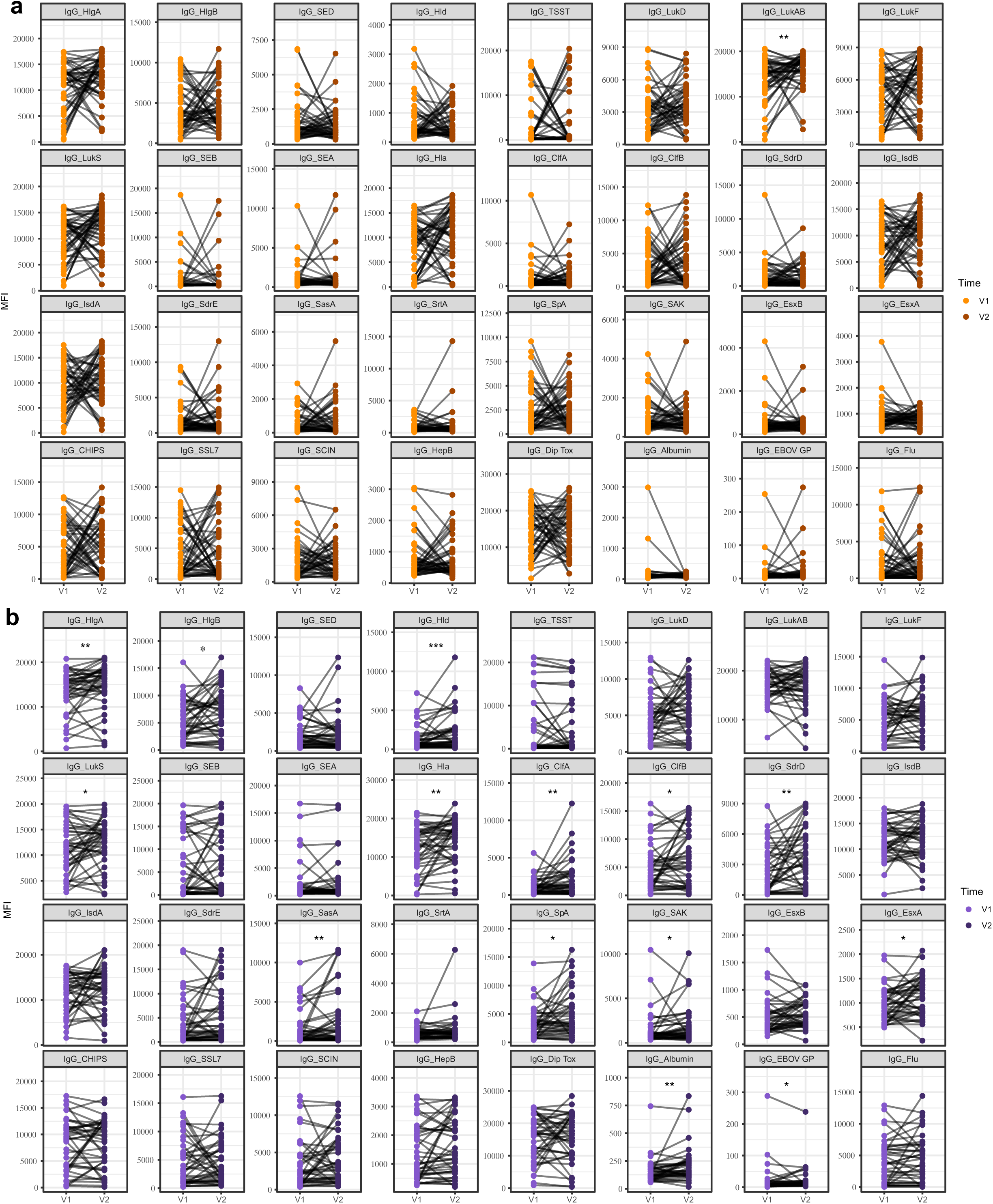
Longitudinal IgG Responses in Pediatric SSTI and Invasive Infections. (a-b) Paired line plots showing antigen-specific IgG responses at acute (V1) and convalescent (V2) timepoints in children with SSTI (a) or invasive (b) infections. Lines connect matched samples from the same individual across timepoints for each antigen feature.

**Figure S5.**
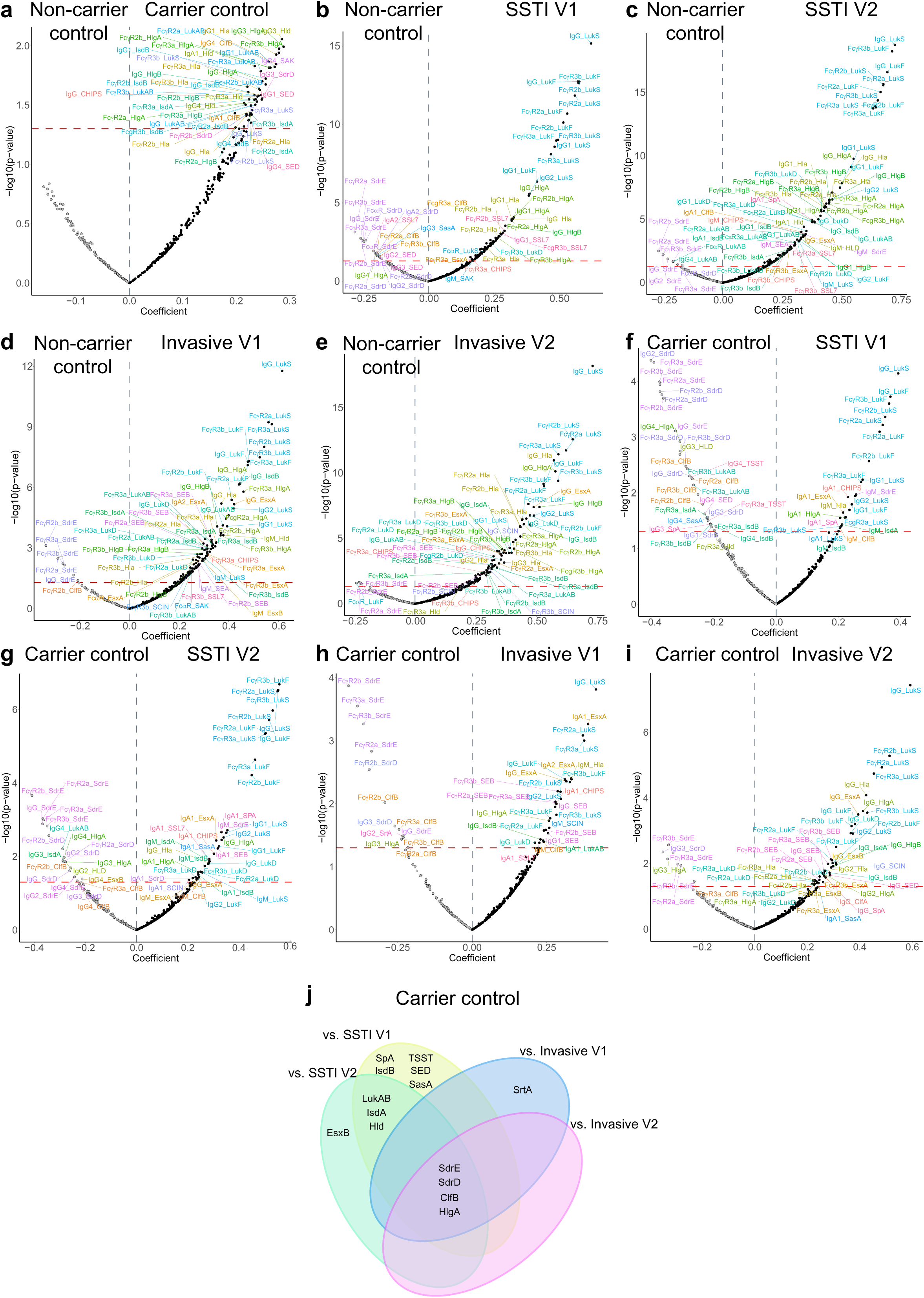
Age-adjusted analyses of humoral responses to *S. aureus.* (a-i) Volcano plots showing antibody features significantly associated with disease state after adjustment for age. Each point represents an antigen–isotype or Fc receptor–binding feature; effect size (x-axis) is the regression coefficient adjusted for age, and significance (y-axis) is the –log₁₀(*p*-value). Features above the horizontal threshold (p <0.05) are significantly enriched in the indicated group. (j) 4-way Venn diagram summarizing features significantly enriched in carrier control individuals compared to disease groups.

**Figure S6.**
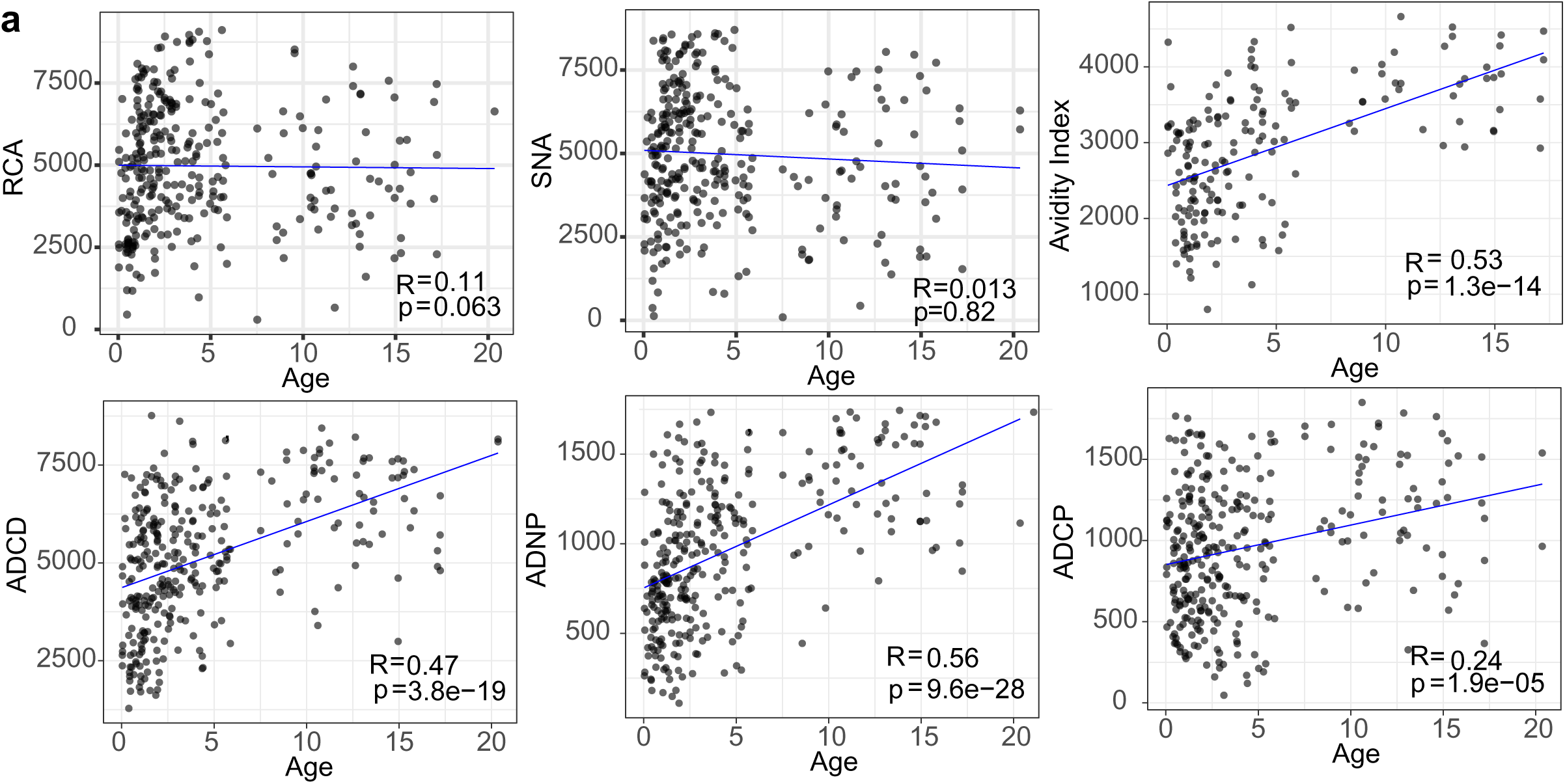
Age-Dependent Maturation of Functional Humoral Responses in Pediatric *S. aureus* Infections. Scatterplots depict the correlation between age (x-axis, in years) and immune features (y-axis, MFI), including lectin binding (SNA), avidity index, and functional antibody activities (ADCD, ADCP, ADNP). Each dot represents an individual subject. Spearman correlation coefficients (R) and corresponding p-values are indicated on each plot.

**Figure S7.**
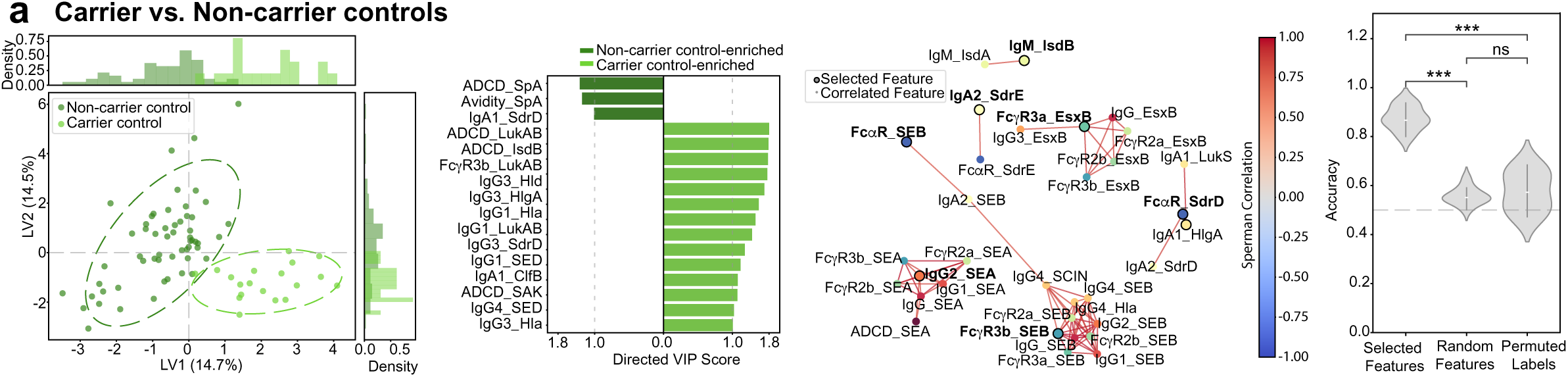
Multivariate modeling distinguishes non-carrier from carrier control. (a) A stability-based LASSO approach was used to identify a minimal set of antibody features that differentiate carrier from non-carrier controls. Age-adjusted antibody measurements were used to construct a partial least squares discriminant analysis (PLS-DA) model. From left to right, panels show: PLS-DA score plots of the first two latent variables demonstrating sample separation; directional bar plots of variable importance in projection (VIP) scores for LASSO-selected features (VIP > 1.0); correlation networks showing co-regulated features based on Spearman correlations (ρ > 0.8, FDR-adjusted p < 0.05); and violin plots of classification accuracy across 100 iterations of 5-fold cross-validation, compared to models built using randomly selected features or permuted class labels.

**Figure S8.**
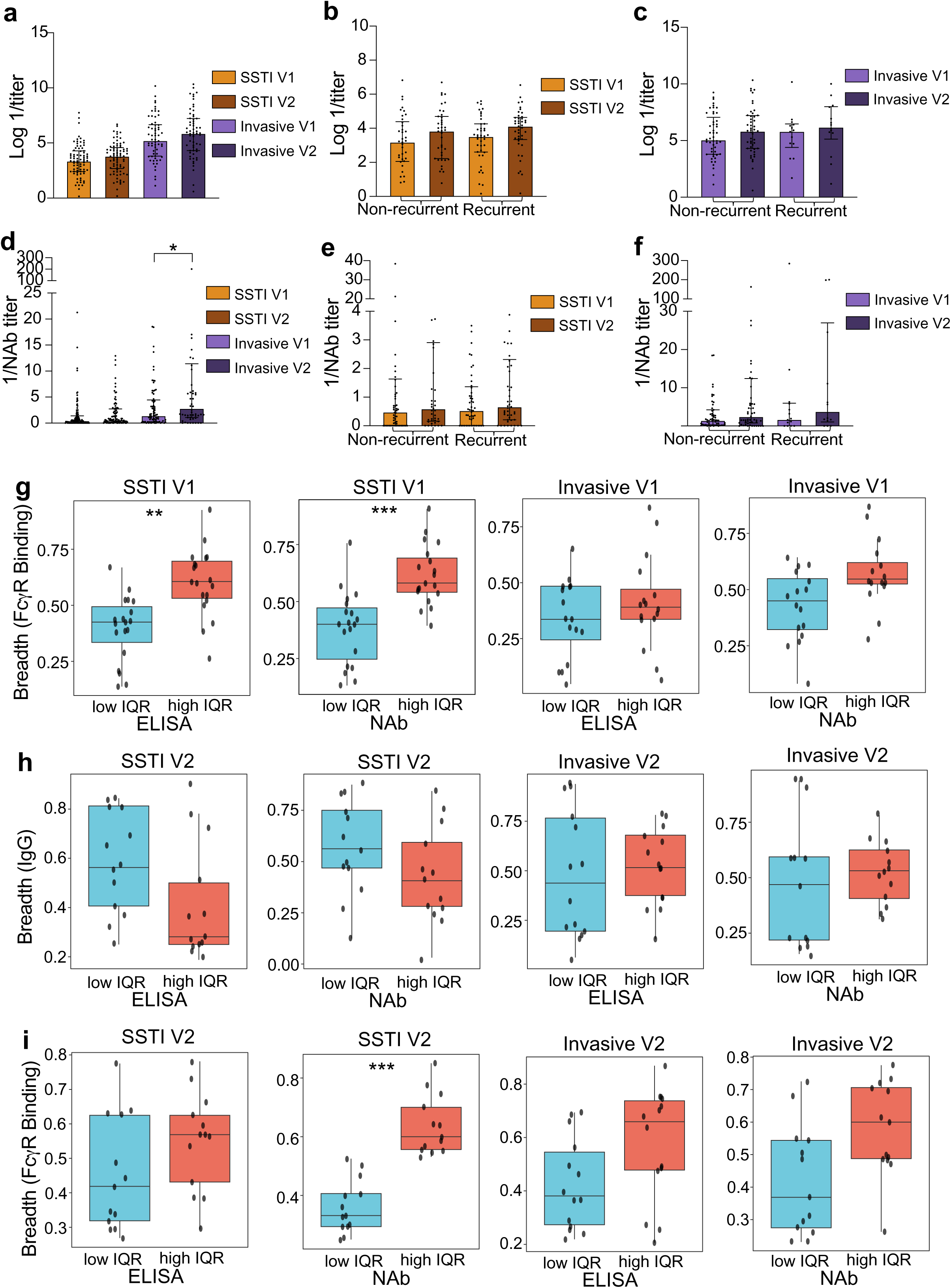
Anti-Hla antibody titers and breadth stratified by recurrence status and disease phase. (a–c) Anti-Hla IgG titers measured by ELISA in children with acute (V1) and convalescent (V2) SSTI and invasive infection, further stratified by non-recurrent versus recurrent disease status. (d–f) Anti-Hla neutralizing antibody (NAb) titers in in children with SSTI and invasive infection, further stratified by recurrence status. (g-i) Antigen breadth of Fcγ receptor–binding or IgG antibody responses in children with SSTI and invasive infection stratified into low- and high-IQR groups based on age-adjusted anti-Hla ELISA or NAb titers. Boxplots indicate the median and interquartile range, with whiskers representing 1.5× IQR; points denote individual subjects. Statistical comparisons between groups were performed using Wilcoxon rank-sum tests with Benjamini–Hochberg correction (**p < 0.05, **p < 0.01, ***p < 0.001).

